# Objective markers of altered neural gain dissociate tinnitus and self-reported listening problems in individuals with exceptional hearing thresholds

**DOI:** 10.64898/2026.10.01.755864

**Authors:** Matthew Paul O’Donohue, Jason Mikiel-Hunter, Heivet Hernández-Peréz, David McAlpine, Juan-Pablo Faúndez-Astudillo

**Affiliations:** Department of Linguistics, The Australian Hearing Hub, Macquarie University, Australia; School of Psychological Sciences, The Australian Hearing Hub, Macquarie University, Australia; Department of Audiology and Speech Pathology. The University of Melbourne, Australia

## Abstract

Many individuals experience problems that are undetected by standard hearing tests, such as persistent tinnitus and difficulties understanding speech in background noise. Here, we report evidence of altered neural gain in young adults with self-reported listening problems despite exceptional hearing thresholds. Individuals reporting persistent tinnitus—the daily percept of sound in the absence of an external source—exhibited elevated thresholds of the middle ear muscle reflex, suggestive of altered gain in brainstem reflexive circuits, and reduced cortical responses to auditory spatial cues conveyed in the temporal fine structure of low-frequency sounds. Moreover, in both groups, greater self-reported difficulty with speech-in-noise listening was associated with higher magnitude cortical responses—particularly at the highest stimulus intensity—consistent with elevated neural gain. Using a conceptual model, we demonstrate how damage to high-threshold auditory nerve fibres innervating cochlear nuclei can account for reduced activity of binaural neurons sensitive to the temporal fine structure of low-frequency sounds, generating a compensatory increase in neural gain in higher brain centres that impacts listening in complex environments.

## Introduction

Complex listening environments present a considerable challenge for the auditory system. Many listeners can process sounds in quiet with relative ease even when their peripheral auditory system is severely damaged (such as cochlear implant and hearing aid users; Lesica, 2018; Wilson & Dorman, 2008). However, this situation changes dramatically in the presence of competing talkers, reverberation, or high levels of background noise. Difficulty understanding speech in noise is one of the most common hearing complaints, even when the audiogram (the standard test of ‘pin-drop’ hearing) suggests that hearing is clinically normal. Problems listening in noise in the absence of elevated hearing thresholds, colloquially referred to as ‘hidden hearing loss’ (HHL; Schaette & McAlpine, 2011), are causing researchers and clinicians to reconsider whether the audiogram is the most appropriate way of diagnosing and assessing the impact of hearing impairment. This is particularly important when the problem for many individuals lies in understanding *speech* in everyday situations that are *not* quiet.

Despite its increasing acceptance as a clinically relevant problem, biomarkers for HHL remain elusive, with no technologies or remediation programs to address it. Neurodegeneration (cochlear synaptopathy), definitively demonstrated in animals (Kujawa & Liberman, 2009), is a likely contributing factor in individuals with HHL. Damage to synapses between sensory hair cells and auditory nerve fibres—as result of exposure to loud sounds and ageing—can spare hearing thresholds even when the damage is substantial (e.g., a loss of 50% of synapses or more; Kujawa & Liberman, 2009, 2015; Oxenham, 2016). However, exploring functional consequences of synaptopathy in humans must contend with the relatively uncontrolled aetiology and nature of occupational and recreational exposure to loud sounds in diverse populations (e.g., musicians, construction workers, nightclub goers), as well as age-related degeneration independent of noise exposure and the likely co-occurrence of audiometric hearing loss (Bharadwaj et al., 2019; Trevino et al., 2023). Conversely, while *in vivo* studies can more easily control for factors such as noise exposure, they are less amenable to exploring subjective experiences such as tinnitus or difficulties communicating in noise. Most *in vivo* studies have explored cellular and neurophysiological processes within the inner ear that are less- or in-accessible *pre-mortem* in humans (Kujawa & Liberman, 2009, 2015). The few studies investigating the consequences of synaptopathy/HHL on downstream processing in the central auditory pathways demonstrate neural substrates at least consistent with subjective human report, including impaired neural representations of speech-in-noise (Monaghan et al., 2020), poorer detection of sounds in noise (Resnik & Polley, 2021), and reduced adaptation to background noise (Bakay et al., 2018). Importantly, these studies report elevated neural gain manifesting as higher spontaneous and evoked firing rates in midbrain neurons (see also Hesse et al., 2016), consistent with a homeostatic increase in central gain following altered sensory input.

The notion that HHL is associated with elevated neural gain was first explored by Schaette and McAlpine (2011), who reported elevated activity in the auditory midbrain in normal-hearing listeners with tinnitus, attributing this to a compensatory response to reduced activity in the auditory nerve. Auditory brainstem responses (ABRs), the compound activity of auditory nerve, brainstem, and midbrain neurons to sound stimulation, represent the most direct measure of neural activity in the human subcortical afferent auditory pathway. The amplitude of wave I of the ABR (generated by the auditory nerve) is reported to be reduced in individuals with persistent tinnitus relative to controls without tinnitus, despite no reduction in the amplitude of wave III (brainstem; Vasilkov et al., 2023) and the later wave V (midbrain; Bramhall et al., 2018; Gu et al., 2012; Kehrle et al., 2008; Schaette & McAlpine, 2011). Nevertheless, as a tool for differential diagnosis of HHL, the ABR suffers from large inter-subject variability (Bharadwaj et al., 2019; Gorga et al., 1988), may be *in*sensitive to loss of high-threshold fibres (Bourien et al., 2014), and is not consistently different between individuals with and without tinnitus (Guest et al., 2017; Milloy et al., 2017; Shim et al., 2017). This has motivated attempts to identify more sensitive measures of synaptopathy/HHL.

Here, we test the hypothesis that an early identifiable manifestation of HHL is altered gain in the central auditory nervous system. To do so, we assessed two objective measures of hearing function, the middle-ear muscle reflex (MEMR) and the interaural phase modulation following response (IPM-FR) in listeners who report tinnitus. Recent evident indicates the MEMR—an efferent reflex primarily driven by high-threshold ANF input (Kobler et al., 1992; Liberman & Klang, 1984)—to be a strong predictor of tinnitus (Vasilkov et al., 2023; Wojtczak et al., 2017) and potentially other factors potentially contributing or related to HHL (older age, high noise exposure, and/or poor speech recognition; Bharadwaj et al., 2022; Shehorn et al., 2020). The IPM-FR—a cortically generated brain potential likely reflecting brainstem activity—is a measure of sensitivity to interaural time differences (ITDs) conveyed in the temporal fine structure (TFS) of low-frequency (<1.5 kHz) sounds. Robust at the individual level, and correlated with perceptual discrimination (Haywood et al., 2015; McAlpine et al., 2016; Undurraga et al., 2016), the IPM-FR is generated by a low-frequency, amplitude-modulated tone that undergoes periodic transitions in interaural phase during amplitude minima in the modulation cycle. The IPM-FR elicits two auditory steady-state responses, an EFR elicited by modulations in the energy envelope (modulated equally in both ears), and a following response (FR) elicited by interaural modulations in the TFS. This paradigm provides several advantages for exploring brain mechanisms contributing to HHL. First, low audiometric frequencies are relatively robust to acoustic injury and the effects of ageing. Any differences in neural responses evident in the IPM-FR are less- or even un-likely to reflect damage, including occult damage, to structures within the inner ear. Second, sensitivity to low-frequency ITD cues is critical to ‘cocktail-party’ listening; different talkers are more distinguishable from each other and from background noise, and speech intelligibility in noise is significantly enhanced (Bronkhorst & Plomp, 1988). Behavioural assays of TFS sensitivity in listeners with normal or near-normal audiometric thresholds show evidence of such impairment (Bernstein & Trahiotis, 2016). Finally, *in vivo* evidence suggests that HHL is associated with elevated central neural gain that extends below the frequency range of the damaging noise (Monaghan et al., 2020). The IPM-FR provides a means of assessing the consequences of elevated central gain at sound frequencies with normal hearing thresholds and potentially unaffected directly by cochlear synaptopathy.

Using this approach, we demonstrate that young, normal-hearing individuals with self-perceived listening problems show evidence of altered central neural gain. Listeners with persistent tinnitus showed elevated MEMR thresholds, consistent with reduced activity in brainstem reflexive circuits controlling the MEMR (Vasilkov et al., 2023; Wojtczak et al., 2017). The magnitude of the TFS component of the IPM-FR, but not that of the energy envelope (EFR), was significantly reduced in listeners with persistent tinnitus. Nevertheless, regardless of tinnitus status, individuals with the most self-reported problems understanding speech in complex listening environments showed the largest magnitude IPM-FRs—consistent with elevated neural gain. This effect was greater at higher compared to lower overall sound intensities. Using a conceptual model, we demonstrate how damage to high-threshold ANFs might lead to a reduction in the activity of ITD-sensitive neurons, generating a compensatory homeostatic increase in neural gain in higher brain centres that impacts listening performance. Neural responses to binaural TFS cues conveyed in low-frequency sounds may provide a particularly sensitive metric to early changes in brain function that impact listening performance before other proposed biomarkers of HHL—elevated EHF thresholds or changes in EFRs—become evident. The dissociable effect of elevated MEMRs thresholds in listeners with tinnitus suggests the possibility of developing a toolbox for differential diagnosis of listening problems in HHL.

## Results

### Young adults with exceptional hearing thresholds and persistent tinnitus show evidence of altered neural gain in a brainstem reflex circuit

To explore neural mechanisms contributing to listening problems absent elevated hearing thresholds, we recruited young adults with persistent unexplained tinnitus (*n* = 12, 19–33 years, *M*_age_ = 25.50 ± 5.18, 9 males) and without (*n* = 16, 18–34 years, *M*_age_ = 21.00 ± 4.00, 6 males). Participants in the tinnitus group had all experienced daily bilateral tinnitus for at least two years, with all participants reporting that it had bothered them in the past year and half reporting a slight or moderate problem with hyperacusis. Upon initial screening, all participants had normal hearing in the standard audiometric range (thresholds ≥ 20 dB HL at 0.25–8 kHz), and unremarkable otoscopy examinations. *Age* (significantly different: *t*(20.10) = 2.50, *p* = .021, *d*_z_ = 0.97) and *sex* (not significantly different: χ^2^(1) = 2.52, *p* = .112) were covariates in all linear mixed models (LMMs) reported here, but results were qualitatively the same in models without these covariates (see *Methods*). Separate LMMs assessing hearing thresholds with *group*, *ear*, and *frequency* as factors showed no effect of *group* in either the standard audiometric range (0.25–8 kHz; *F*(1, 24) = 0.92, *p* = .348) or in the extended high-frequency (EHF) range (10–16 kHz; *F*(1, 24) = 0.07, *p* = .799, Fig. 1a). Similarly, an LMM performed on distortion product otoacoustic emissions (DPOAEs), which index the function of outer hair cells, using *group, ear,* and *frequency* (12 levels: f_2_ = 0.5–10 kHz) as factors, also showed no effect of *group* (*F*(1, 24.03) = 1.16, *p* = .293, Fig. 1b). There were also no interactions involving *group* in any of these analyses (9 tests, all *p* > .060).

**Fig. 1.**
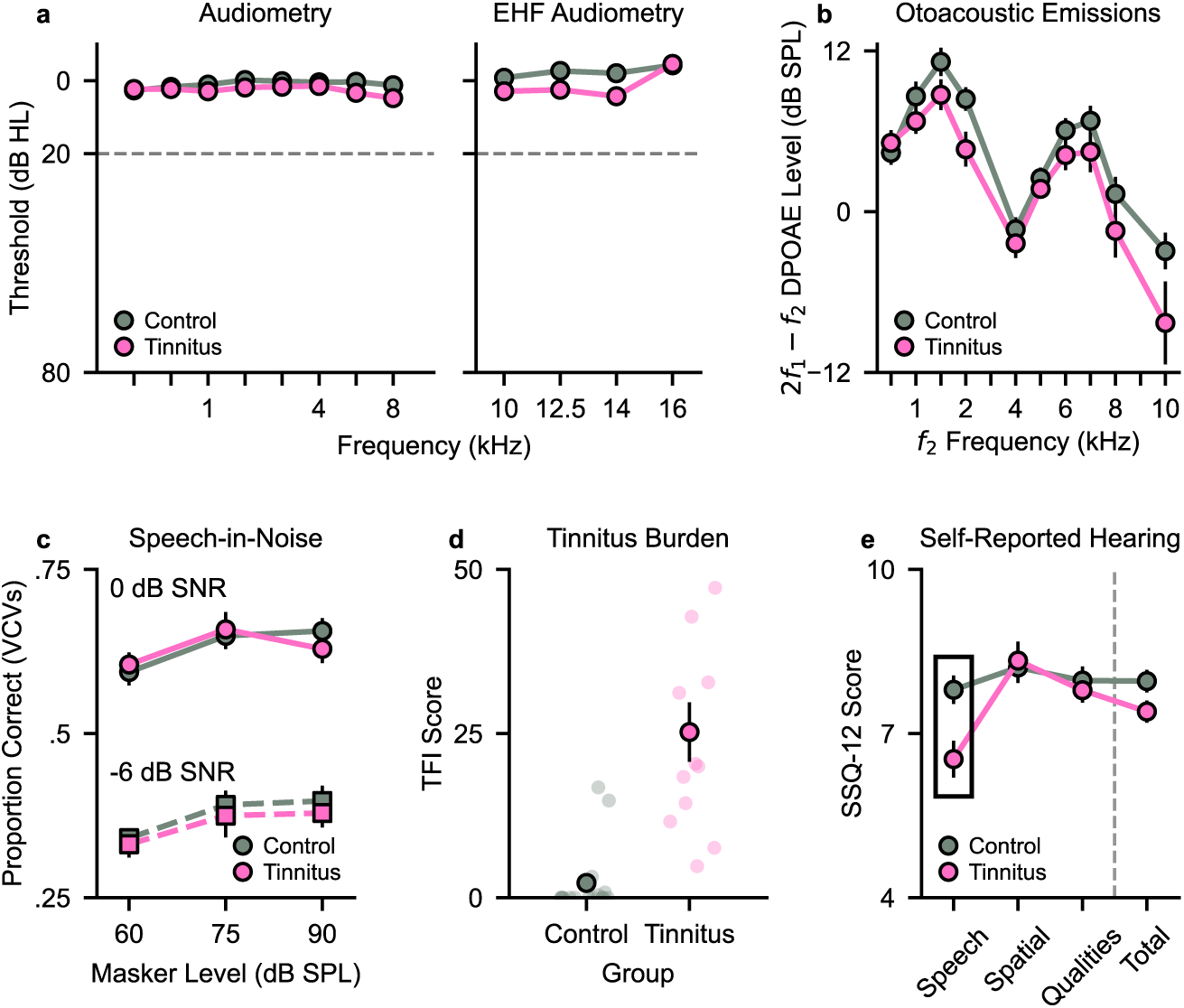
Measures of peripheral and subjective auditory functioning. There were no significant differences between control (grey) and tinnitus (pink) groups in mean **a** low frequency (left panel) and extended high frequency (EHF, right panel) audiometric thresholds for warble tones (thresholds above dashed line at 20 dB HL are considered clinically normal), **b** distortion product otoacoustic emission (DPOAE) 2f2−f1 levels, and **c** proportion of consonants identified in vowel-consonant-vowel tokens presented in 16-talker babble (a test of speech-in-noise perception; SNR is signal-to-noise ratio). **d** Scores on the Tinnitus Functional Index (TFI) survey, which assesses the functional impact of tinnitus in the past week, indicated considerable variation in tinnitus burden despite all participants with tinnitus having persistent (i.e., daily) symptoms. Two controls had modest TFI scores despite reporting no tinnitus on the screening survey (ESIT-Q), but results did not depend on whether these participants were included in the control group. **e** The tinnitus group had a significantly lower mean score on the SSQ-12 Speech subscale (highlighted by the surrounding box), indicating they had greater self-reported difficulty with listening to speech in background noise. In all panels, error bars denote ±1 standard error of the mean (SEM; not visible in **a** due to the y-axis scale).

We next assessed speech-in-noise perception by presenting vowel-consonant-vowel (VCV) tokens in 16-talker babble through insert earphones, with listeners using a computer mouse to click on the correct consonant. An LMM testing *group*, *signal-to-noise ratio* (-6, 0 dB SPL), and *masker level* (60, 75, 90 dB SPL) on VCV discrimination showed no effect or interaction involving *group* (main effect: *F*(1, 24) = 0.59, *p* = .450; interactions all *p* > .524, Fig. 1c). Nevertheless, our subjective measure of listening difficulties—the 12-item version of the Speech, Spatial, and Qualities of Hearing Scale (SSQ-12)—revealed lower SSQ-12 Speech scores in listeners with tinnitus relative to the control group (Welch’s *t*(22.27) = -2.97, *p* = .007, *d*_z_ = -1.14). Thus, our sample of listeners with tinnitus reported listening difficulties (persistent tinnitus, Fig. 1d, and speech-in-noise problems, Fig. 1e) despite having exceptional hearing thresholds, normal DPOAE levels, and normal speech-in-noise (VCV) performance—all of which indicate healthy auditory functioning by clinical standards.

Recent studies of normal-hearing listeners suggest the middle-ear muscle reflex (MEMR) is a reliable indicator of a tinnitus percept (Vasilkov et al., 2023; Wojtczak et al., 2017). This is consistent with reduced efferent feedback from high-threshold auditory nerve fibres— implicated in cochlear synaptopathy—innervating the stapedial muscle of the middle ear. We assessed the strength of the MEMR in our tinnitus and control groups and found that listeners with tinnitus showed significantly weaker MEMRs. Ipsilaterally-evoked MEMRs were recorded using two different probe tones (226 Hz, 1000 Hz), and a wideband noise elicitor that increased in level in 3 dB HL steps from 65 to 98 dB (Fig. 2a). This provided MEMR growth-with-level data for each probe tone and ear in just a few minutes of testing (Fig. 2b). We fitted 3-parameter sigmoid functions to participant data and modelled the 50% threshold (Fig. 2c) in an LMM including factors *group* and *probe tone*, finding an elevated threshold in tinnitus relative to controls (main effect *group*: *F*(1, 22.60) = 5.34, *p* = .030), with a larger effect for the 1000-Hz probe tone (*interaction*: *F*(1, 23.08) = 12.42, *p* = .002). This was supported by an LMM on the raw growth-function data (with *level* now a factor) showing significant interactions of *group* x *level* (*F*(11, 1126.35) = 2.90, *p* < .001), *group* x *probe tone* (*F*(1, 1131.39) = 31.53, *p* < .001), and *group* x *level* x *probe tone* (*F*(11, 1126.35) = 2.63, *p* = .003). We considered whether the weaker MEMR in the tinnitus group might be accounted for by their lower SSQ-12 Speech scores but found that these were not significantly associated with MEMR thresholds at the bivariate level (controls: all *p* > .658, tinnitus: all *p* > .144) and in an LMM (no main effect of *speech* or interaction with *group* or *probe tone*, all *F* < .32, all *p* > .578). These data support previous findings of a weaker MEMR in tinnitus (Vasilkov et al., 2023; Wojtczak et al., 2017), hypothesised to result from cochlear neural degeneration that reduces afferent input into brainstem reflexive circuits innervating the stapedial muscle, the main effector of the MEMR.

**Fig. 2.**
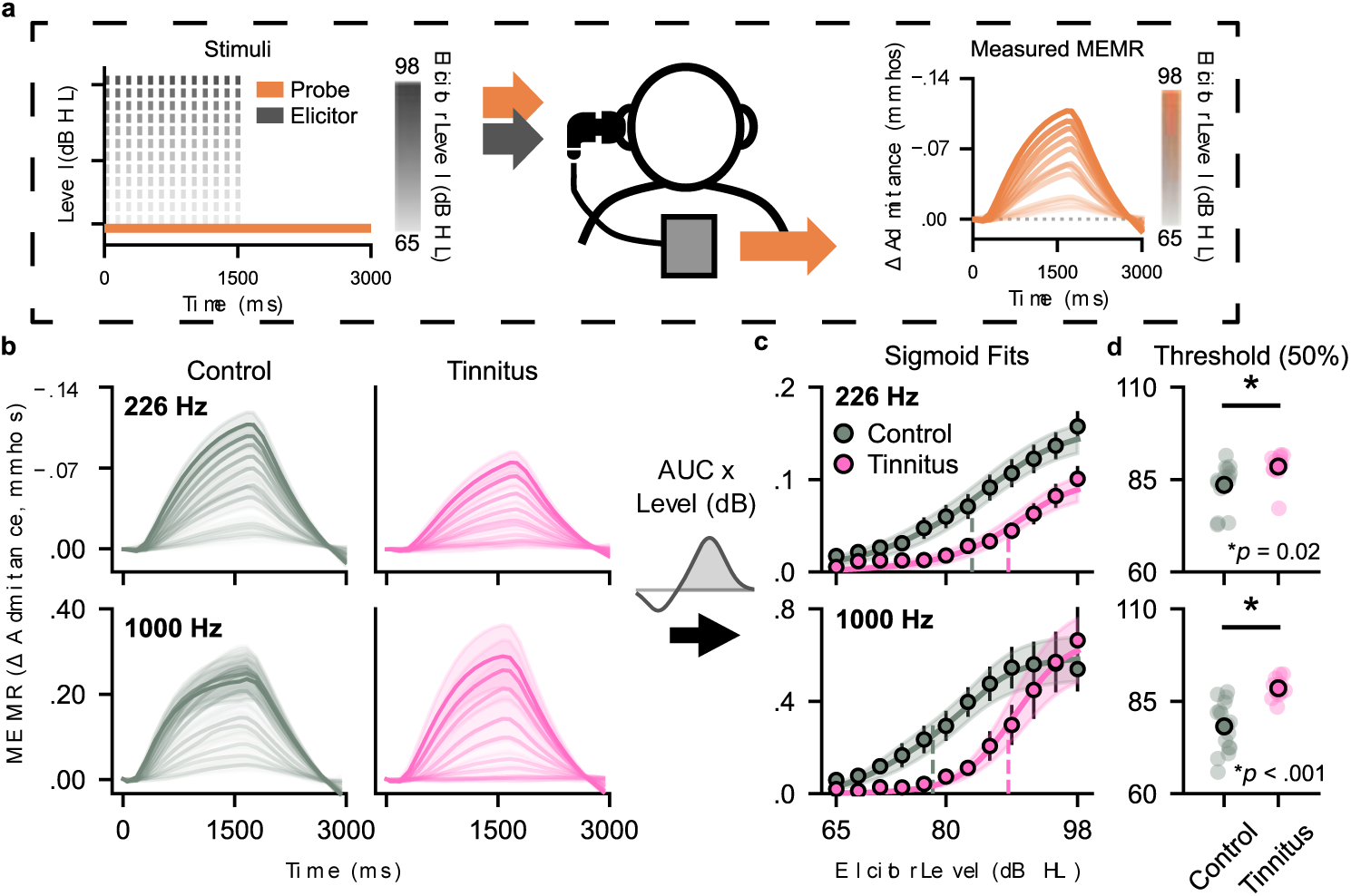
Middle ear muscle reflex measurements. **a** Schematic showing how 226 Hz MEMRs were obtained using Interacoustics Titan Suite settings. The 226-Hz probe tone (orange line) was presented at 85 dB SPL for the full 3-s duration of each run, while a wideband elicitor (grayscale tiles) was pulsed 12 times within approx. the first 1500 ms (70 ms interstimulus interval). Across runs, the level of the elicitor increased from 65 to 98 dB HL in 3 dB steps. The probe tone and elicitors were delivered into the same ear (i.e., an ipsilateral MEMR configuration), with MEMR strength measured from this same ear as the change in admittance evoked by the elicitor. The procedure for the 1000-Hz probe was similar, but the elicitor was presented 13 times per run. **b** Mean MEMRs (averaged across two ears) for probe tones 226 Hz and 1000 Hz show the growth in MEMRs with increasing elicitor level. For each probe tone and elicitor level, the area under the curve for each probe’s canonical response (negative/descending for 226 Hz, positive/ascending for 1000 Hz) was used as the mean MEMR depicted in **c**, where solid lines show three-parameter sigmoid functions fitted to each participant’s data by minimising the residual sum of squares. These “area under the curve” values were highly positively correlated with the maximum MEMR value or the “mean of area” value generated by the Titan Suite software, demonstrating that our results do not depend on the method of measurement (see *Supplementary Information* Fig. S1). **d** Mean 50% thresholds, obtained from these fitted sigmoids, were significantly higher in tinnitus relative to controls (*p* values are from Welch’s *t*-tests). Raw MEMR-by-level data and fitted sigmoid functions for each participant are provided in *Supplementary Information* Fig. S2. In all panels, shaded regions and error bars denote ±1 SEM (not visible in **d** due to y-axis scale).

### Listeners experiencing permanent tinnitus show reduced magnitude binaural cortical-evoked responses

Tinnitus and noise-induced synaptopathy have both been associated with aberrant (elevated) neural activity in the central auditory pathways. To assess changes in central auditory processing associated with tinnitus and perceived listening problems, we measured electroencephalography (EEG) responses while participants listened to a 500-Hz tone amplitude-modulated at 40 Hz (evoking an envelope following response, EFR) with 7-Hz interaural phase difference (IPD) transitions (evoking an interaural phase modulation following response, IPM-FR, Fig. 3a–d). Stimuli were presented at 65 dB(A) and 80 dB(A), in two configurations of IPD transition: -90°↔90° and 0°↔180°. We hypothesised that changes in neural activity linked to low-frequency ITD processing, which is critical to speech perception in noisy environments and which depends on temporally precise inputs that are likely sensitive to neural gain changes, would be associated with self-reported speech-in-noise difficulties (a core symptom of HHL). This is indeed what we observed. At a bivariate level, elevated IPM-FR amplitudes in the -90°↔90° condition were significantly and strongly associated with *poorer* SSQ-12 Speech scores in the tinnitus group for the 80 dB stimulus (tinnitus *r* = -0.73, *p* = .007; control *r* = -0.38, *p* = .141) and to a marginal degree for the 65 dB stimulus (tinnitus *r* = -0.57, *p* = .052; control *r* = -0.11, *p* = .697, Fig. 3f). An LMM testing *group*, *level*, and (SSQ) *speech* showed significant effects of *level* (IPM-FR larger at 80 dB than 65 dB; *F*(1, 24) = 15.68, *p* < .001), *group* (IPM-FR lower in tinnitus group than in control group; *F*(1,22) = 5.39, *p* = .030), and *speech* (IPM-FR larger with lower *speech* scores; *F*(1, 22) = 6.16, *p* = .021) and significant interactions of *level* x *speech* (*F*(1, 25) = 5.92, *p* = .023) and *group* x *level* (*F*(1, 24) = 5.66, *p* = .026). The *level* x *speech* interaction shows that the *speech* x IPM-FR association was stronger at 80 dB, the higher stimulus level (*p* = .003), than at 65 dB (*p* = .235). Equivalently, this shows that lower *speech* scores were associated with greater IPM-FR amplitude growth from 65 to 80 dB, suggesting that changes in IPM-FR within an individual participant (i.e., a self-normalising measure) may be predictive of speech-in-noise difficulties (Fig. 3g). The lack of a *group* x *speech* or *group* x *speech* x *level* interaction (*F*(1, 24) < .24, *p* > .632) does not support the view that this relationship depends on having tinnitus, although it was descriptively more robust in the tinnitus group at the bivariate level (which does not adjust for age and sex; Fig. 3f). Notably, the *group* x *level* interaction resulted from IPM-FR amplitudes in the tinnitus group being significantly lower than controls at 80 dB (*p* = .005) but not 65 dB (*p* = .224, Fig. 3e).

**Fig. 3.**
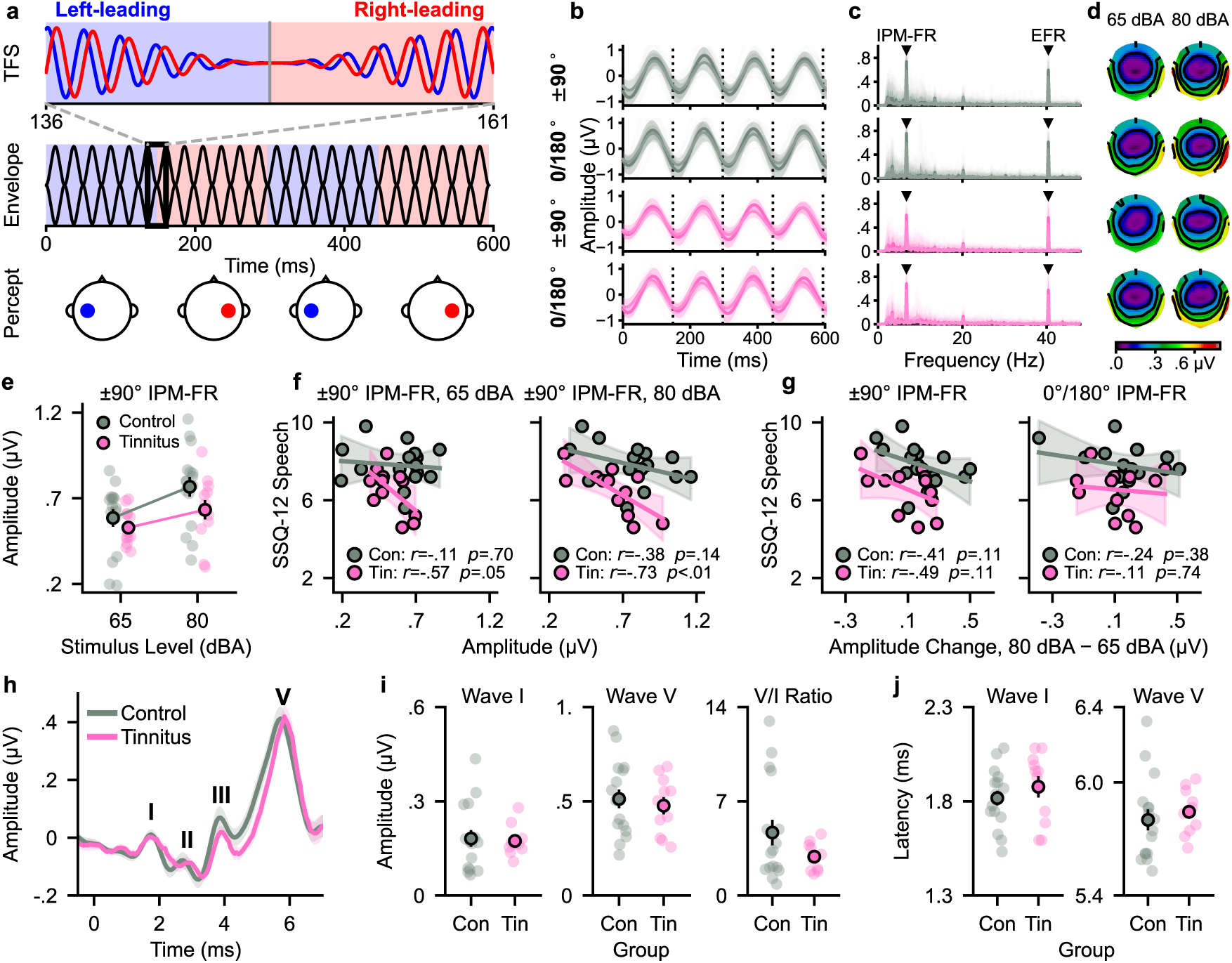
Neural markers of elevated gain. **a** IPM-FR stimulus schematic, with top panel depicting a −90° to 90° IPD transition embedded in the TFS (temporal fine structure), middle panel depicting the envelope modulated at ∼40 Hz, and blue and red lines indicating 500-Hz tones presented to left and right ears, respectively, with colour shading to indicate the IPD as left-leading (blue) or right-leading (red). Bottom panel depicts the lateralised sound image for a given IPD. **b-d** EEG responses in **b** time and **c** frequency domains and **d** topographic maps. Top two rows depict control group responses for the −90°↔90° (denoted ±90°) and 0°↔180° (denoted 0/180°) IPD conditions, and bottom two rows depict tinnitus group responses for these conditions. Stimulus level is represented as different lines in time- and frequency-domain panels (65 dBA: light, thin; 80 dBA: dark, thick) and as different columns for topographic maps. Time- and frequency-domain data were taken from the sensor with the largest amplitude across all participants (M2), with standard deviation shown by the shaded area. In the time domain, vertical dotted lines show IPD transition times. In the frequency domain, faint lines shown individual responses and triangular markers indicate the frequency of the IPM-FR (∼7 Hz) and EFR (∼40 Hz). **e** Mean −90°↔90° IPM-FR amplitudes (measured in the frequency domain and averaged across M1 and M2) were lower for the tinnitus group than for controls at the higher stimulus level (80 dBA), and **f** higher amplitudes were associated with lower SSQ-12 Speech scores, again at the higher stimulus level only (shown in right panel; left panel shows 65 dBA stimulus). This indicates elevated neural activity in individuals who report greater difficulty listening to speech in noisy environments. The LMM did not provide evidence that this relationship was specific to the tinnitus group (Tin), despite the bivariate scatterplots here (which do not adjust for age and sex) suggesting a more robust relationship in tinnitus than in controls (Con). **g** This level-dependent effect can also be expressed as a significant negative association between SSQ Speech scores and the growth in −90°↔90° IPM-FR amplitudes from 65 to 80 dBA (left panel), which was not observed for the 0°↔180° stimulus (right panel). Thus, IPM-FR amplitude changes across stimulus levels may have potential as a self-normalising measure within individual participants. **h** Mean (and ±1 SEM shaded region) ABRs to an 11 Hz click train, with waves I, II, III, and V labelled. For each participant, we averaged electrode channels ipsilateral to the ear of stimulus delivery. **i** Mean ABR wave I and V amplitudes and **j** latencies were not significantly different between control (Con) and tinnitus (Tin) groups. In all panels, error bars denote ±1 SEM.

These data suggest two independent gain mechanisms operating within our sample: a general reduction in IPM-FR amplitudes in the tinnitus group, consistent with a mechanism that disrupts the fine temporal processing required for tracking ITDs, and an increase in IPM-FR amplitudes in individuals with poorer self-reported speech listening (regardless of tinnitus prevalence), consistent with evidence that auditory neural hyperactivity reduces the neural discriminability of speech at high intensities (Monaghan et al., 2020). Both these effects were observed at the higher, but not lower, overall intensities. Equivalent models for the EFR (which does not require binaural integration of sub-millisecond ITDs) and the 0°↔180° IPM-FR (a binaural test based on differential excitation/inhibition) conditions did not reveal any association between EEG amplitudes and *group* or *speech* (*p* > .188 for all main effects and interactions involving these factors), with significant main effects only for *level* (amplitudes larger at 80 dB compared to 65 dB; all *p* < .004). Scatterplots depicting (non-significant) bivariate associations between SSQ Speech scores and EFR and 0°↔180° IPM-FR amplitudes are provided in Supplementary Fig. S3.

Evidence for a role of elevated central gain in tinnitus was initially based on the relative amplitudes of waves I and V in the ABR, but the inter-subject variability in the ABR and its dependence on demographic variables such as sex (Bharadwaj et al., 2019) suggests alternative measures, such as the IPM-FR, are required. To assess whether central gain changes were also evident in the ABR here, we calculated wave I and V peak-to-trough amplitudes from monoaurally evoked, ipsilaterally-recorded responses to broadband clicks, and calculated the ratio of wave V to wave I from each participant’s ABR traces (averaged across mastoid electrodes M1 and M2; Fig. 3h). These data were then modelled with an LMM including factors *group* and *channel* (M1, M2). While amplitudes and ratios were in a similar range to those reported by Schaette and McAlpine (2011), there were no significant effects of *group* or *group* x *channel* interactions (all *p* > .165; Fig. 3i), although this is perhaps unsurprising given the mixed sexes and young ages in our (small) samples; indeed, the only significant effect was the expected main effect of covariate *sex* on wave V amplitude (smaller in males; *p* = .039). Similar results were obtained in LMMs modelling wave I and V latencies (longer wave V latency in males, *p* < .001; all other *p* > .109, Fig. 3j).

### Identifying early markers for the trajectory of HHL symptoms

Identifying early biomarkers of HHL has clinical relevance, providing a means of tracking, and potentially intervening in, the development of hearing problems. To this end, we explored whether our objective measures and self-reported listening difficulties could reveal a trajectory of symptoms by stratifying our sample into four subgroups: 10 listeners from our control group with no HHL symptoms (CON+), the other 6 listeners from our control group, who explicitly reported at least a slight “difficulty listening to speech in a noisy situation” on the ESIT-Q (*n* = 4) and/or TFI scores indicating some tinnitus in the past week (despite reporting “no tinnitus” on the ESIT-Q; *n* = 2), 5 listeners not included in the tinnitus group (and therefore not included in any previous analyses) due to having less-than-daily tinnitus (TIN−), and our 12 listeners with persistent (i.e., daily) tinnitus (TIN+). The subgroups with intermediate HHL symptoms (CON+ and TIN−) exhibited an intermediate MEMR profile (Fig. 4a), with growth functions larger than the persistent tinnitus group (TIN+) but smaller than controls with no HHL symptoms (CON−). This was tested by modelling MEMR thresholds obtained from the fitted sigmoid functions in an LMM including fixed factors *group* (CON−, CON+, TIN−, TIN+) and *probe tone.* We found significant effects of *group* (*F*(3, 25.76) = 4.59, *p* = .010) and *probe tone* (*F*(1, 26.28) = 18.46, *p* < .001), and a significant interaction *F*(3, 26.22) = 4.48, *p* = .012). *Post hoc* tests for the 1000-Hz probe, where the effect of *group* was most robust, showed elevated thresholds in TIN− (*p*_Holm_ = .006) and TIN+ (*p*_Holm_ < .001), and possibly CON− (*p*_Unc_ = .026 non-significant after Holm correction; *p*_Holm_ = .106), relative to CON+ (Fig. 4b).

**Fig. 4.**
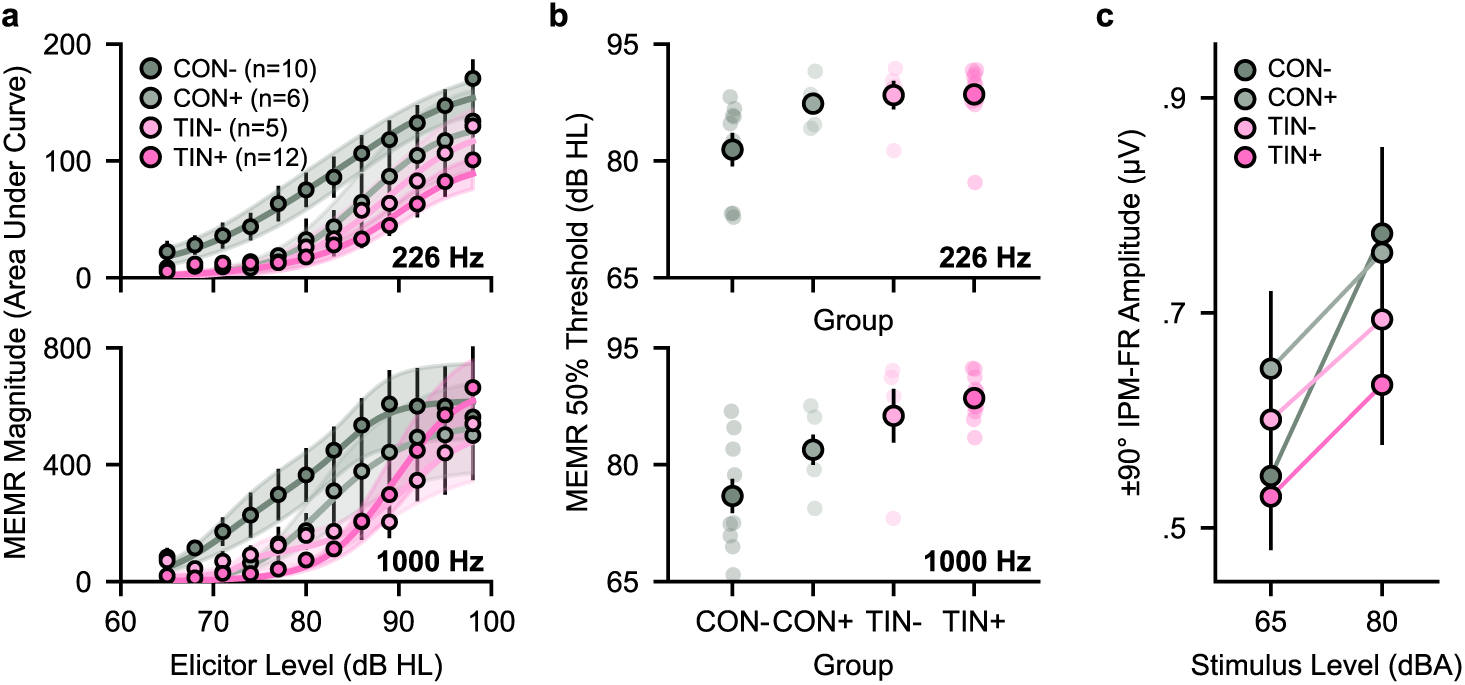
Subgroup analyses suggest a trajectory of HHL symptoms. **a** Sigmoid functions fitted to MEMR growth-by-level data and **b** 50% thresholds suggest a weakening MEMR with increasing HHL symptoms (speech-in-noise difficulties and tinnitus), although the small samples warrant caution in interpretation. **c** Mean −90°↔90° IPM-FR amplitudes were variable across subgroups but remained consistent with lower amplitudes in participants with tinnitus at the higher stimulus level. In all panels, error bars and shaded regions denote ±1 SEM. CON−, controls with no HHL symptoms; CON+, controls who indicated some speech-in-noise difficulties and/or recent tinnitus; TIN−, less-than-daily tinnitus; TIN+, daily tinnitus.

MEMR thresholds between the three groups with HHL symptoms were not significantly different (all *p*_Unc_ > .087, *p*_Holm_ > .25), but plots depict a trajectory of weakening MEMRs with increasing symptoms of HHL, with thresholds being highest—and most consistently high—for participants with persistent tinnitus. This was supported by post-hoc polynomial contrasts that revealed a significant linear trend across the groups for MEMR thresholds at 1000 Hz (*t*(34.6) = 4.48, *p* < .001), although linear and quadratic terms for 226 Hz MEMR thresholds did not reach significance (linear *t*(36.3) = 1.76, *p* = .087; quadratic *t*(38.4) = -1.82, *p* = .077). Despite the caution associated with a small sample size, these data support the potential utility of the MEMR for early diagnosis of HHL at the individual level. We also tested whether IPM-FR amplitudes followed an HHL trajectory across the four subgroups. As in the main two-group analysis, lower SSQ Speech scores were associated with larger amplitudes at 80 dB (*speech* × *level* interaction, *p* = .025; no main effect of *speech*, *p* = .111) and there was a significant *group* × *level* interaction (*p* = .034). This pattern matched the two-group result, with TIN+ having lower amplitudes than controls at 85 dB (vs CON−: *p* = .030; vs CON+: *p* = .060, both uncorrected), although no pairwise contrast survived Holm correction (all *p*_Holm_ > .182; Fig. 4c). Post-hoc polynomial contrasts showed a significant linear trend across the groups for IPM-FR amplitudes at 80 dB (CON− < CON+ < TIN− < TIN+, *t*(28.5) = 2.43, *p* = .022), but it is worth noting that IPM-FR amplitudes were quite variable in these small subgroups. Overall, intermediate HHL symptoms were associated with higher MEMR thresholds relative to symptom-free controls, with some indication that this trajectory was also associated with reduced IPM-FR amplitudes at the higher stimulus level.

## Discussion

Employing standard clinical hearing tests, subjective listening assessments, and two objective measures of brain function, we find that young individuals reporting persistent tinnitus, but exceptional hearing thresholds show significantly higher MEMR thresholds and significantly lower cortical-evoked (IPM-FR) responses compared to controls. Moreover, greater IPM-FR responses were associated with greater self-reported speech-in-noise listening difficulties (as assessed with the SSQ-12) in both groups. Importantly, only the magnitude of the binaural (TFS) component of the IPM-FR—which tracks the interaural cue conveyed in the sub-millisecond TFS and not the component tracking the modulation of sound energy (the EFR)— was reduced in the tinnitus group and associated with SSQ-12 scores. Our data support the interpretation that a reduced magnitude MEMR is a hallmark of those with tinnitus and a normal audiogram (Vasilkov et al., 2023; Wojtczak et al., 2017). Our data also support the contention that binaural cues conveyed in the TFS of low-frequency sounds provide for a sensitive marker of listening problems in those with normal hearing thresholds.

### Altered neural gain is associated with self-reported hearing problems

In addition to tinnitus, a critical feature of HHL seems to be (generally self-reported) problems listening in high levels of background noise. Here, we found that listening difficulties were associated with larger IPM-FR magnitudes. At the bivariate level, this relationship was only significant for the tinnitus group at the higher sound level (i.e., at 80 dB, not 65 dB), but the full model (adjusting for age and sex) suggested it was present in both groups (and more robust at the higher level). These results suggest that a cortical marker of altered gain is characteristic of self-perceived speech-in-noise problems, potentially independent of tinnitus, which is more evident at higher than at lower sound intensities. This is consistent with *in vivo* models of HHL that are characterised by impaired sound intensity adaptation (Bakay et al., 2018), and less discriminable neural representations of human speech sounds (vowel-consonant-vowel) in noise (Monaghan et al., 2020), at high sound intensities.

### A conceptual model of elevated neural gain can explain cortical responses that are reduced in tinnitus and larger in listeners with speech-in-noise difficulties

High-threshold auditory-nerve fibres (ANFs) are most susceptible to damage from transient exposure to high intensity sounds (though low- and medium-threshold fibres might also be implicated; Encina-Llamas et al., 2019) and contribute to the gain of the middle ear through a feedback circuit innervating the stapedial muscle (Kobler et al., 1992; Liberman & Klang, 1984). A weaker MEMR (i.e., a higher MEMR threshold and/or shallower MEMR growth with level) has been reported in normal-hearing listeners who report tinnitus (Vasilkov et al., 2023; Wojtczak et al., 2017) and in mice with cochlear synaptopathy following noise exposure (Valero et al., 2016, 2018). Nevertheless, a tinnitus percept is unlikely to arise from reduced gain in the MEMR circuit *per se*. Rather, tinnitus is likely a consequence of compensatory elevated gain in the central auditory nervous system following reduced sensory input, including through temporary ear-plugging (Brotherton et al., 2019; Hutchison et al., 2023).

Here, we describe a conceptual model wherein reduced input from high-threshold ANFs elicits a reduction in feedback gain to the middle ear muscles but an increase in feedforward gain that disrupts neural processing of auditory spatial cues at multiple stages within the central auditory pathways (Fig. 5). Importantly, our model shows how elevated feedforward gain in the early brainstem, rather than the disrupted ANF function that reduces gain in the MEMR circuit, can account for reduced cortical responses to the TFS of low-frequency sounds.

**Fig. 5.**
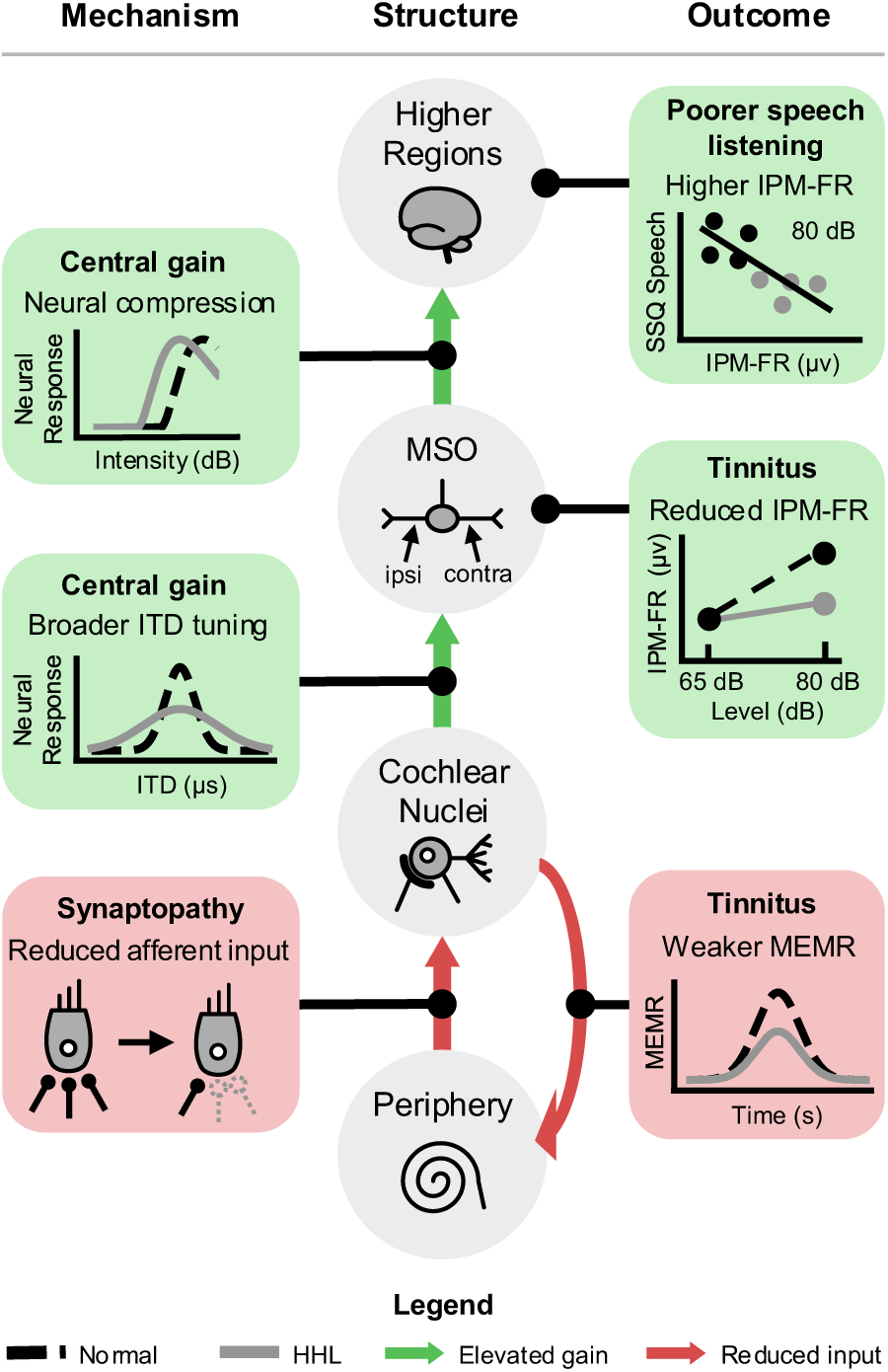
A conceptual model of altered neural gain in HHL. Boxes on the left describe proposed mechanisms of gain-related changes in auditory processing, circles in the middle represent auditory pathway stages, and boxes on the right represent outcomes of altered gain as measured in the current study. In the model, synaptopathy reduces neural output from the cochlea in the auditory periphery, which reduces the efferent activity that drives the middle ear muscle reflex. A compensatory increase in neural gain disrupts the precise timing of binaural inputs to the MSO (medial superior olive), resulting in reduced IPM-FR amplitudes in individuals with tinnitus at the higher stimulus level. Elevations in neural gain in higher brain structures (e.g., midbrain, thalamus, cortex), which might induce neural compression (e.g., rollover) at lower stimulus intensities, result in difficulties listening to speech in noisy environments irrespective of tinnitus, as evidenced by the negative correlation between SSQ-12 Speech scores and IPM-FR amplitudes at the higher stimulus level.

In the model, action potentials phase-locked to the TFS of low-frequency sounds are conveyed from each ear by ANFs innervating neurons in the anterior division of the ventral cochlear nucleus (AVCN) and then to primary binaural neurons of the medial superior olive (MSO) in left and right brainstem (Grothe et al., 2010). MSO neurons, acting as binaural coincidence detectors, respond maximally when action potentials from each ear arrive in close temporal alignment. Sources of internal noise that jitter this binaural alignment reduce maximum binaural output, similar to how reduced interaural coherence—the extent to which sound arriving at the two ears is identical—reduces performance in localisation tasks (Faller & Merimaa, 2004), and makes sounds, including speech, harder to detect (Luke et al., 2022) or understand in background noise (Lavandier & Culling, 2010). Cochlear synaptopathy could introduce internal noise that jitters binaural alignment by disrupting phase locking of action potentials in the auditory nerve or by reducing activity in high-threshold ANFs, which elicits a compensatory increase in brainstem excitability that renders binaural temporal integration less precise.

We consider the latter explanation more likely in listeners with tinnitus. Low- and medium-threshold ANFs, rather than the thin and poorly myelinated high-threshold ANFs, provide the primary input to the specialised AVCN neurons subserving temporally precise ITD processing; damage to high-threshold ANFs directly is unlikely to impair phase-locking in brainstem binaural pathways. Further, even if synaptopathy extended to low- or medium-threshold ANFs (e.g., Encina-Llamas et al., 2019), the frequency at which we elicit the IPM-FR—500 Hz—lies below the range most associated with noise-induced hearing loss. Other than reduced MEMRs, our listeners with tinnitus show no other evidence of cochlear dysfunction, with comparable hearing thresholds to controls extending into the EHF range, as well as present and comparable DPOAEs, suggesting only subtle deficits in nerve function are likely. Notably, *in vivo* studies suggest elevated neural gain extending below the region of temporarily damaging noise is a hallmark of HHL (Monaghan et al., 2020), which supports rate coding for auditory features but may disrupt precise temporal processing (Chambers, Resnik, et al., 2016). For these reasons, we favour an explanation in which reduced activity in high-threshold ANFs at frequencies higher than 500 Hz leads to broadband elevated neural gain (see also Noreña, 2011) in brainstem pathways, including those subserving binaural hearing.

A similar increase in neural gain—occurring after binaural integration—could explain the relationship between subjective hearing problems and the cortically-evoked IPM-FR. A wealth of neurophysiological evidence has demonstrated neural hyperexcitability or disinhibition following cochlear synaptopathy in auditory midbrain, thalamus, and cortex (Asokan et al., 2018; Balaram et al., 2019; Chambers, Resnik, et al., 2016; Chambers, Salazar, et al., 2016; Eggermont, 2017; Hesse et al., 2016). This has been linked to degraded neural representations (Chambers, Resnik, et al., 2016; Monaghan et al., 2020) and poorer behavioural detection (Resnik & Polley, 2021) of sounds in background noise. This explanation also makes sense from the perspective of the well-documented rollover effect, whereby speech-in-noise understanding increases with increasing intensity before falling as intensities exceed approximately 85–90 dB (Jerger & Jerger, 1967, 1971), including in normal-hearing listeners (Hornsby et al., 2005; Studebaker et al., 1999). The roll-over effect is also linked to degraded neural representations (Horvath & Lesica, 2011; Wong et al., 1998). A cogent explanation for listening problems associated with higher magnitude IPM-FRs, therefore, is that elevated central gain pushes listeners into rollover at relatively lower overall sound intensities. This is akin to neurophysiological evidence in gerbils that amplification in the hearing aid improves audibility of quiet speech but exacerbates difficulties listening to speech at high intensities by inducing rollover at lower-than-normal intensities (Armstrong et al., 2022). Indeed, the relationship between self-reported listening and IPM-FR magnitudes in our data was more robust for the 80 dB stimulus than the 65 dB stimulus, nearer to the sound intensities at which rollover occurs in normal-hearing listeners. Note that such a listening impairment can arise in the absence of elevated hearing thresholds and without the need to invoke a direct role for high-threshold ANFs in the neural coding of sounds in high levels of background noise (Carney, 2018). For example, the latter has been proposed as an explanation for reduced EFR magnitudes in individuals with tinnitus (Paul et al., 2017) and poorer suprathreshold hearing (Bharadwaj et al., 2015; Mepani et al., 2021)—a pattern not observed in the current study— but this view has been contested (Encina-Llamas et al., 2019).

Our simple model proposes that persistent unexplained tinnitus is characterised by cochlear synaptopathy that reduces afferent input into the MEMR circuit, and a compensatory increase in neural gain (before binaural integration) that reduces activity in MSO neurons that code ITDs conveyed in the TFS of low-frequency sounds. Then, regardless of tinnitus prevalence, elevated gain following binaural integration leads to self-perceived listening deficits, possibly by reducing neural dynamic range—an effect that becomes most pronounced at high overall sound intensities, where neural firing rates are more likely to reach saturation, leading to a more compressed neural representation of sound. We suggest our model may provide a framework for future investigations into the precise gain mechanisms that are implicated in HHL.

### Identifying HHL in individual listeners

Understanding hearing problems in individuals with normal audiometric thresholds, and translating findings to clinical settings, is challenging given the breadth of factors that can explain an individual’s symptoms, including middle- and inner-ear pathologies and demographic factors such as age and sex (Bharadwaj et al., 2019). Moreover, existing measures of neural gain changes in HHL, such as the auditory brainstem response (Gu et al., 2012; Schaette & McAlpine, 2011; Vasilkov et al., 2023), show large variability between individuals (Bharadwaj et al., 2019; Gorga et al., 1988) and inconsistent support as a biomarker for HHL (Guest et al., 2017; Milloy et al., 2017; Shim et al., 2017), including in our data here (although this is perhaps unsurprising since our sample was small, young, and had both sexes; cf. Gu et al., 2012; Schaette & McAlpine, 2011). We therefore aimed to test a potentially sensitive electrophysiological biomarker of HHL—evoked by a stimulus highly relevant for speech-in-noise understanding—while tightly controlling various peripheral, demographic, and other factors (e.g., somatic or vascular causes of tinnitus) that could confound group effects. This resulted in a small sample (12 with tinnitus and 16 without) which, though warranting caution in interpreting our results, provides for strict control over sample characteristics—a key feature of our approach. Our data suggest that when peripheral auditory functioning (included extended high frequency audiometric thresholds) and likely non-synaptopathic causes of tinnitus are controlled for, biomarkers of HHL may be evident in individuals of a very young age—well before age-related degeneration of peripheral or auditory nerve structures or the onset of confounding sensorineural hearing loss.

Our data are consistent with a weakening MEMR as listeners accumulate symptoms of HHL. While individuals with persistent tinnitus had consistently high MEMR thresholds relative to individuals with no HHL symptoms (who showed lower and more variable MEMR thresholds), individuals with intermediate HHL symptoms—those explicitly reporting at least a slight speech-in-noise listening difficulty and/or intermittent tinnitus—had MEMR thresholds that were consistently between these groups. Previous studies have also observed a weakened MEMR in participant groups with suspected cochlear synaptopathy (Bharadwaj et al., 2022; Vasilkov et al., 2023; Wojtczak et al., 2017), at least when tonal elicitors are not used (Guest et al., 2019). Thus, our results lend further support to the MEMR as an early biomarker of HHL even in small samples, including when using a setup common in audiology clinics.

While we could not identify such a trajectory using the IPM-FR, our data suggest that level-dependent growth in this measure was associated with self-reported listening problems. On average, individuals in the tinnitus group exhibited little-to-no growth in the IPM-FR as stimulus level increased (from 65 dB to 80 dB), while controls exhibited greater IPM-FR amplitudes at the higher intensity. Furthermore, in both groups, individuals with poorer self-reported listening exhibited greater IPM-FR amplitudes at the higher intensity. Thus, while inter-subject variability may create difficulties in interpreting the IPM-FR when it is measured at a single stimulus intensity, measuring the IPM-FR across intensities (or potentially other features such as the magnitude of the interaural phase differences) may have clinical utility, especially when combined with other measures such as the MEMR. Such an approach is also highly feasible; the IPM-FR can be obtained with just five minutes of recording and with methods similar to those currently used in clinical settings (e.g., for ABRs). However, future studies, particularly those with larger samples, are needed to verify that the MEMR and the IPM-FR are sensitive markers of HHL at the individual participant level.

## Conclusion

Our data suggest that alterations in neural gain at multiple stages in the central auditory pathway can explain problems undetected by standard hearing tests. Listeners with persistent unexplained tinnitus exhibited a weaker MEMR, suggestive of reduced afferent input to the cochlear nucleus, as well as reduced cortical responses on a measure of low-frequency binaural hearing (the IPM-FR), consistent with a compensatory increase in neural gain that disrupts precise temporal coding in the MSO. Regardless of tinnitus status, however, greater IPM-FR magnitudes were associated with self-reported difficulties processing speech in noisy or complex listening environments, consistent with elevated gain in higher auditory brain centres. Our key measures implemented here—the SSQ-12, the MEMR, and the IPM-FR—are quick to administer and leverage standard clinical methodologies; together, they may have potential for detecting hearing problems not evident in the audiogram or standard electrophysiological measures of auditory brainstem function.

## Acknowledgements

This research was supported by Tinnitus UK, the Martin Lee Centre for Innovations in Hearing Health, and the Macquarie University Research Acceleration Scheme. We thank Jessica Monaghan and Lina Yao for support with funding, Jaime Undurraga for developing the pEEGy package and stimulus delivery tools, and Jayde Cahir for support with project coordination.

## Author Contributions

Conceptualization, DM, JPFA, JMH, MO, HHP; Data Curation, MO, JPFA, JMH; Formal Analysis: MO, JPFA, JMH, DM; Funding Acquisition, DM, JMH, JPFA; Investigation, MO, JPFA; Methodology, JPFA, MO, JMH, HHP; Project Administration, MO, HHP, JPFA, DM; Resources, MO, JPFA, JMH; Software, MO, JPFA, JMH, HHP; Supervision, MO, HHP, DM, JPFA; Validation, MO, JPFA, DM, JMH, HHP; Visualization, MO, JPFA, DM, JMH, HHP; Writing – Original Draft, MO, DM; Writing – Review & Editing, MO, DM, JPFA, HHP, JMH

## Data and Code Availability

All data and code used for analyses will be made publicly available at Figshare upon publication: https://doi.org/ 10.25949/34021098.

## Methods

### Participants

Participants were recruited via adverts disseminated through social media, Macquarie University noticeboards, and professional networks such as local hearing clinics. Participants volunteered for the experiment, gave written informed consent, and were compensated A$20/hr for their time. The study was approved by the Human Research Ethics Committee of Macquarie University (ref: 520261075366932).

### Procedure

#### Subjective and Behavioural Measures

Before being invited to participate in the experiment, prospective participants were screened remotely using the European School for Interdisciplinary Tinnitus Research Screening Questionnaire (ESIT-Q) to ensure they did not have a hearing loss diagnosis, a (cochleo-)vestibular dysfunction,or an obvious peripheral, somatic, vascular, and/or otherwise objective cause of tinnitus (e.g., pulsatile tinnitus, temporomandibular joint dysfunction, **history** indicative of Meniere’s Disease). The experiment was conducted in an electromagnetically and acoustically isolated room at the Australian Hearing Hub on Macquarie University campus. First, otoscopy was performed on both ears and participants with excessive earwax or an abnormal tympanic membrane were excluded. Second, audiometric thresholds were measured using the Interacoustics Affinity Suite (Compact Version 4) and a warble tone stimulus presented via RadioEar DD450 headphones to each ear at the following frequencies: 0.25, 0.5, 1, 2, 3, 4, 6, 8, 10, 12.5, 14, and 16 kHz. Participants were excluded if any thresholds in the 0.25–8 kHz range (obtained using the modified Hughson Westlake descending method) were above 25 dB HL. Third, tympanometry, middle ear muscle reflex (MEMR), and distortion product otoacoustic emission (DPOAE) measurements were obtained in each ear using the Interacoustics Titan Suite. Tympanometry and MEMRs were measured using probe tones of 226 and 1000 Hz. Elicitor stimuli for MEMRs were wideband noises (0.4–12 kHz) presented from 65–98 dB HL in 3 dB steps (Fig. 2a). Using the Titan Suite default settings, the MEMR measurement at each elicitor level took 3 s, with the probe tone presented throughout the duration and 12 (226 Hz probe) or 13 (1000 Hz probe) elicitor presentations throughout the first half of each measurement (70 ms interstimulus intervals resulting in approx. 1500 ms duration). Measurements were repeated if an artifact was observed. DPOAEs were obtained with an 2f_2_/f_1_ frequency ratio across an f_2_ sweep of 0.5–10kHz across 12 frequencies at 65 (f_1_) and 55 (f_2_) dB SPL.

Next, participants completed a speech-in-noise perception task on a computer delivering using MATLAB R2019b. Vowel-consonant-vowel (VCV) stimuli were presented in sixteen-talker babble delivered diotically through ER-3C insert earphones. Following each trial, participants used a mouse to click on the consonant in the VCV target, with the next trial starting automatically after the response. There were six blocked and randomly ordered conditions consisting of two target-to-masker ratios (0 and -6 dB SPL) and three masker levels (60, 75, and 90 dB SPL), with 42 trials per condition.

Then, during EEG setup (below), participants completed three surveys delivered through the website REDCap: the Speech, Spatial and Qualities of Hearing Scale (12-item version; SSQ-12), assessing self-reported hearing across three subscales, and the Tinnitus Functional Index (TFI), assessing the functional impact of tinnitus in the past week.

### EEG

EEG recordings were obtained using a BioSemi ActiveTwo system, with 64 scalp electrodes (placed in accordance with the international 10-20 system), two mastoid electrodes (left/M1 and right/M2), and two extraocular electrodes, placed on the and left lateral canthus and right infraorbital area . Electrode voltage offsets were usually below 20 mV and never exceeded 40 mV. The sampling rate of recordings was 16384 Hz for all participants except for one participant (see Exclusions). The EEG session began with 5 mins of open-eyes resting state followed by 5 mins of closed-eyes resting state (for a related study), with participants instructed to sit as still as possible and minimise eye movements. Next, we recorded ABRs by presenting 100-µs clicks at 100 dB p–peSPL through ER-3C insert earphones connected to an RME Fireface UC sound card (24 bits, 48-kHz sampling rate). Clicks had alternating polarity, a mean rate of 11 or 40 Hz (with presentation timing jittered uniform randomly in a range spanning ±5% of the interstimulus interval), and were presented monoaurally to each ear separately, resulting in four randomly ordered blocks (2 rates × 2 ears) of 4000 repetitions each. Participants were instructed to keep their eyes closed throughout the recordings.

For the IPM-FR recordings, stimuli were ∼498.5 Hz carrier tones amplitude modulated at ∼40.4 Hz and with an interaural phase difference (IPD) that periodically and symmetrically alternated at ∼6.7 Hz (Fig. 3a). These carrier and modulation frequencies ensured an integer number of carrier and amplitude modulation cycles within each IPD cycle. In each of four randomly ordered 5-min blocks, stimuli were presented at either 65 or 80 dB(A) and with an IPD modulation between 0° and 180° or between -90° and 90°. Each IPD modulation was applied at a minimum in the amplitude modulation cycle to prevent monoaural cues in the stimulus. Stimuli were delivered through ER-3A insert earphones. During the recordings, participants watched a silent, subtitled nature documentary.

### Statistical Analyses

#### Data Exclusions and Cleaning

One tinnitus participant was excluded from MEMR analyses for both probe tones, and two controls for the 226 Hz probe tone only, due to unusual raw MEMR responses or poor sigmoid fits to MEMR-by-level data. Including them did not affect results. Two of these participants had non-standard (Type A) tympanograms due to an admittance peak slightly above or below admittance limits (i.e.,Types As and Ad); two others (one control, one tinnitus) also had non-standard tympanograms, but excluding all four participants from MEMR and ABR analyses did not affect results. One datapoint for the 226 Hz MEMR analysis was imputed via linear interpolation due to an extreme and atypical response (see Supplementary Fig. S2). One tinnitus participant was missing from ABR analyses due to an incorrect EEG sampling rate of 4096 Hz.

#### Statistical Models

Linear mixed models (LMM) were fit with the afex package in R-4.5.1 using the Satterthwaite approximation for degrees of freedom. All LMMs included full factorial terms for fixed factors (e.g., *group* x *probe tone* x *level*), age and sex as covariates, and participant as a random intercept without random slopes as these structures did not converge reliably. We used age as a covariate instead of hearing thresholds since these two variables are strongly associated (Mepani et al., 2020) and since only age was significantly different between the control and tinnitus groups. All code for analyses is available online (see *Data and Code Availability*).

#### EEG Analyses

For pre-processing EEG data, we used Python 3.10 packages MNE 1.10.1 (for ABRs) and pEEGy 1.5.9 (for the IPM-FR; https://gitlab.com/open-source-brain/peegy). For ABR analyses, recordings were referenced to the vertex electrode Cz, bandpass filtered (100–1500 Hz), and averaged across 4000 epochs per condition. Waveforms were time-shifted forward by 1 ms to correct for acoustic delay introduced by insert earphones. Amplitudes for Waves I, III, and V were manually marked from peak to following trough by two researchers (one with clinical experience) who were blinded to any participant or group information.

For IPM-FR analyses, recordings were referenced to Cz and downsampled to 1024 samples per second. Poor electrode contacts (typically 1 or 2 electrodes with large DC components) and eye blinks were automatically detected and removed (see Valderrama et al., 2018). Data were then filtered between 2 Hz and 60 Hz for frequency analyses, and between 2 Hz and 8 Hz for time-domain plots. Data were filtered using a finite impulse response Kaiser filter (60 dB ripple in pass and stop bands, 1 Hz transition width) with the filter’s delay compensated to give non-causal zero-phase filtering. Data were then divided into 68 epochs of ∼4.45 s each (providing 30 amplitude modulation cycles per epoch) and de-noised using denoising source separation (DSS; (De Cheveigné & Simon, 2008). Denoised epochs were weighted averaged and then the frequency response was computed (fast Fourier transform of 4560 points at ∼0.22 Hz resolution) and the IPD and amplitude modulation rates extracted as spectral components for statistical analyses.

