## Supplementary Figures 1-3 for "Objective markers of altered neural gain dissociate tinnitus and self-reported listening problems in individuals with exceptional hearing thresholds"

**Supplementary Materials**

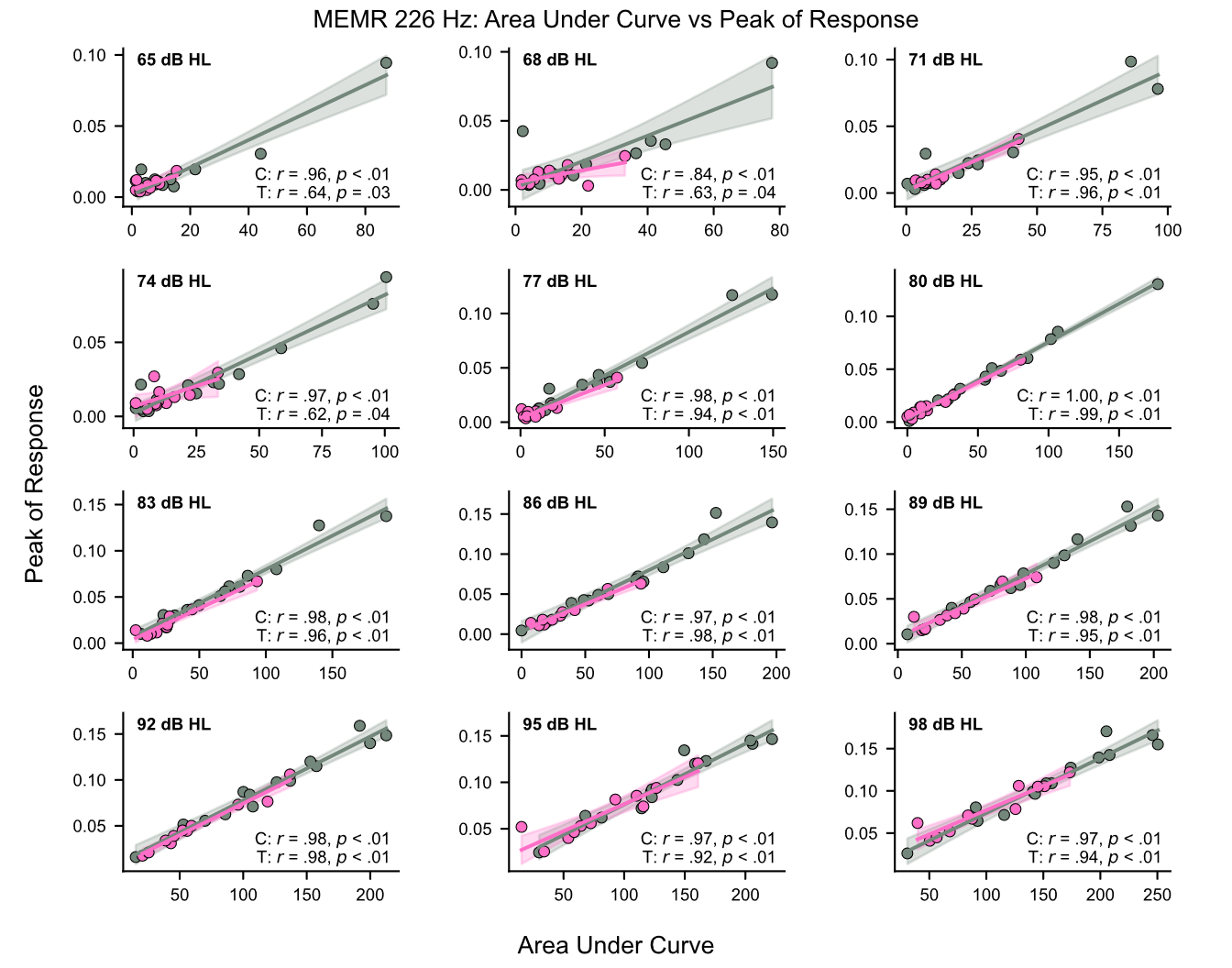

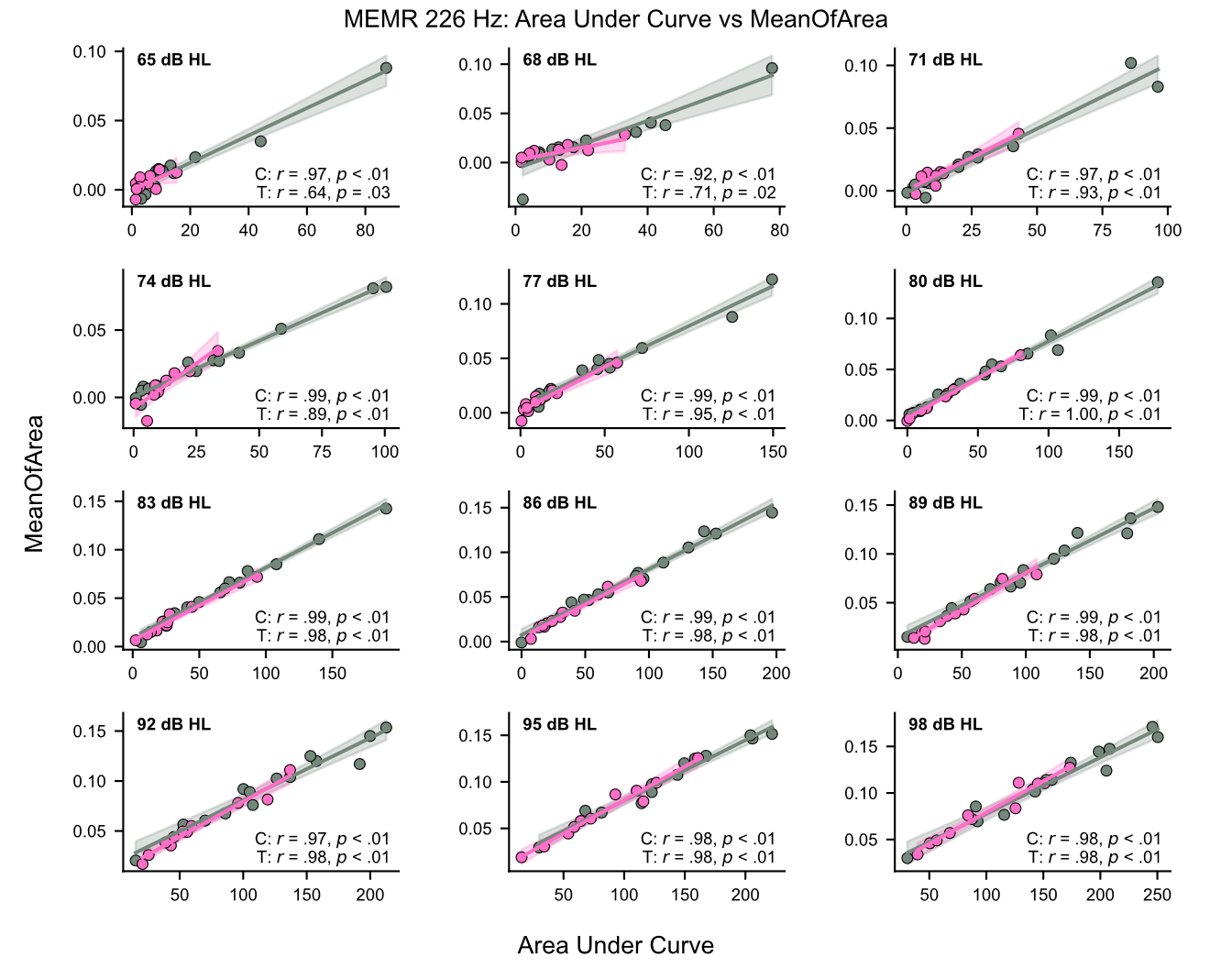

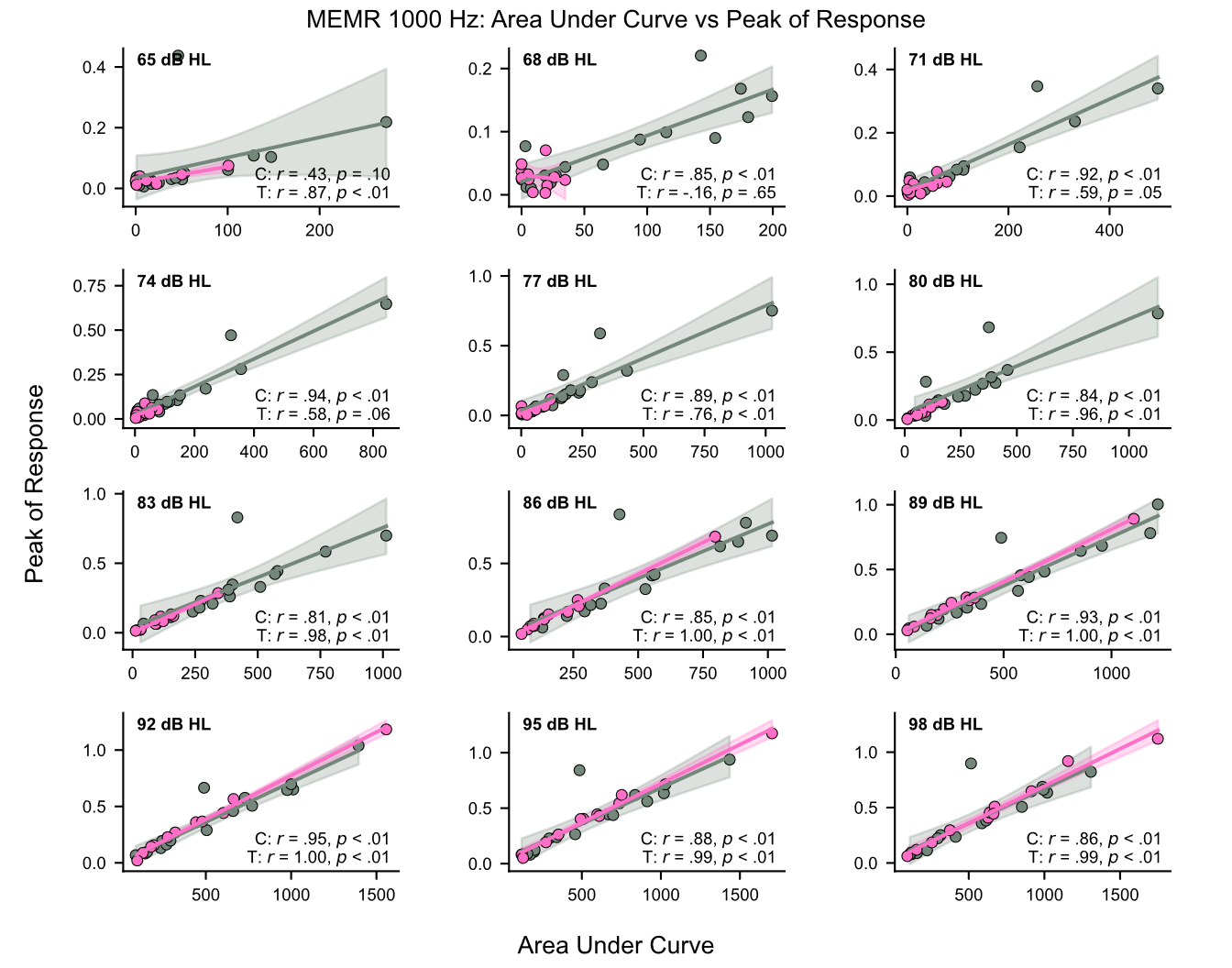

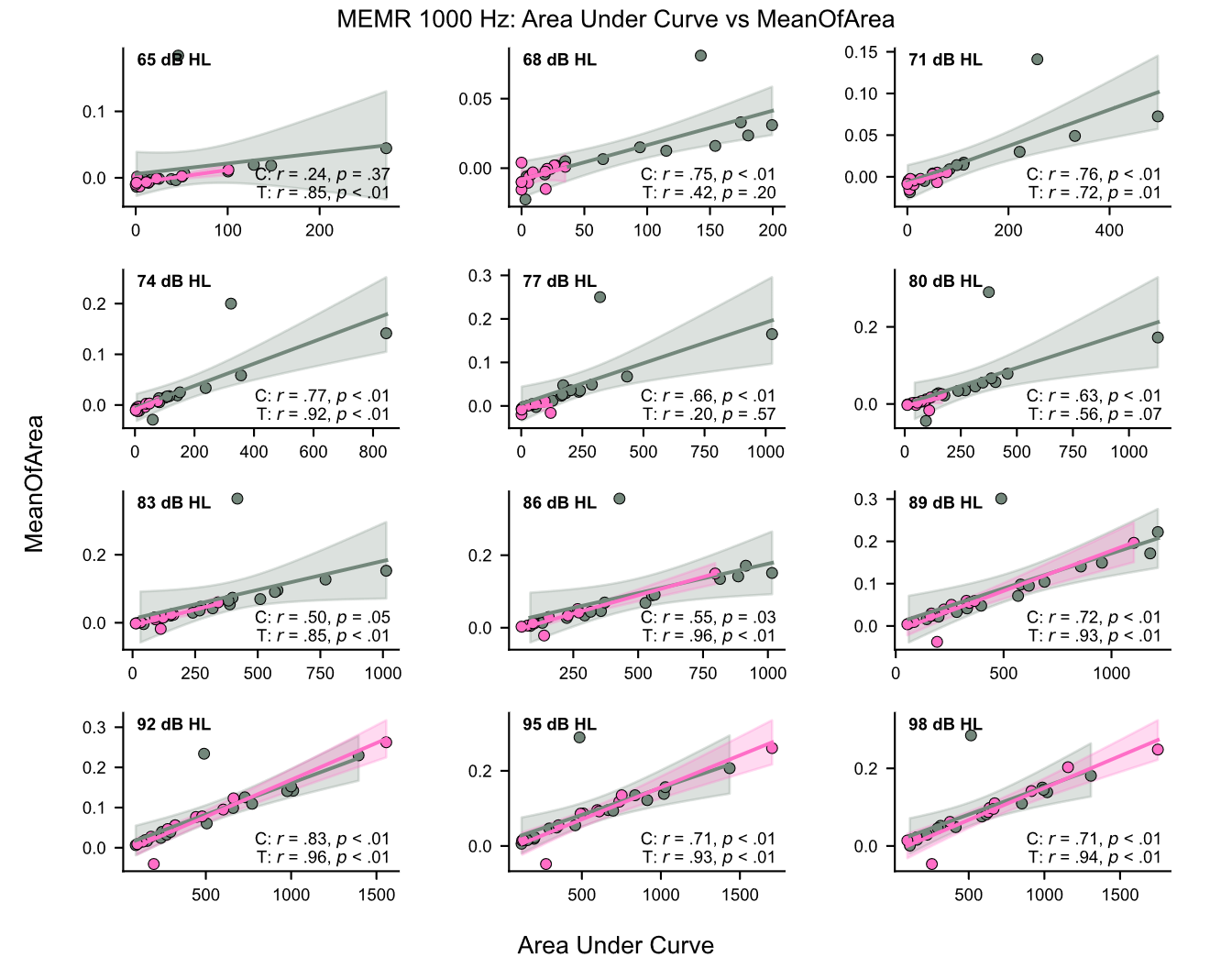

**SUPPLEMENTARY FIGURE S1.** Bivariate scatterplots comparing methods for obtaining the middle ear muscle reflex (MEMR) magnitude at each elicitor level (shaded regions show the 95% confidence interval of the linear fit). In the main text, MEMR values were the area under the curve of the MEMR; in both the control and tinnitus groups, these values were strongly positively correlated with the maximum value of the MEMR (“peak of response”) and the “meanOfArea” value provided by the Interacoustics Titan Suite, suggesting that our MEMR threshold estimation was not method dependent. C, control group; T, tinnitus group.

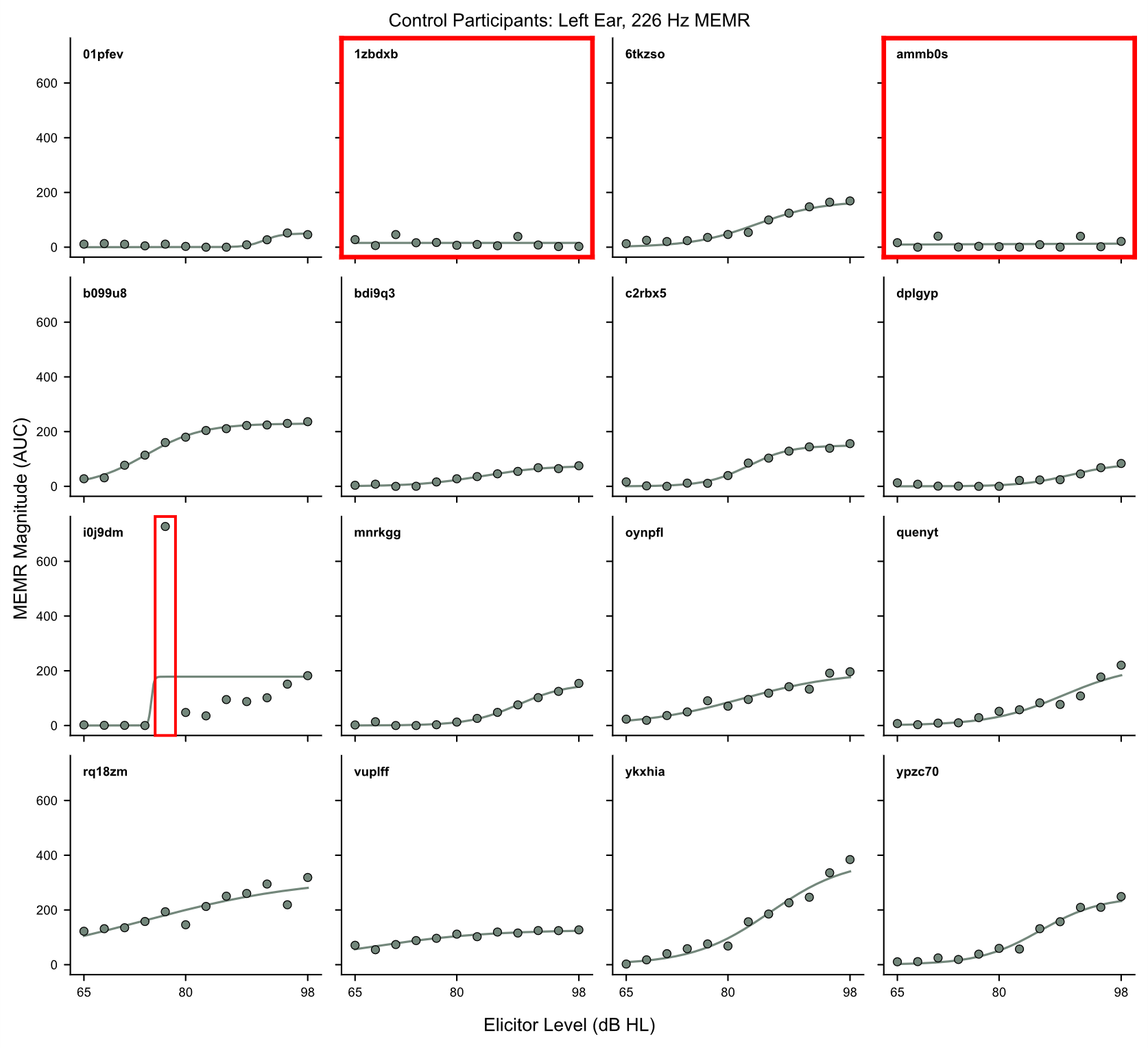

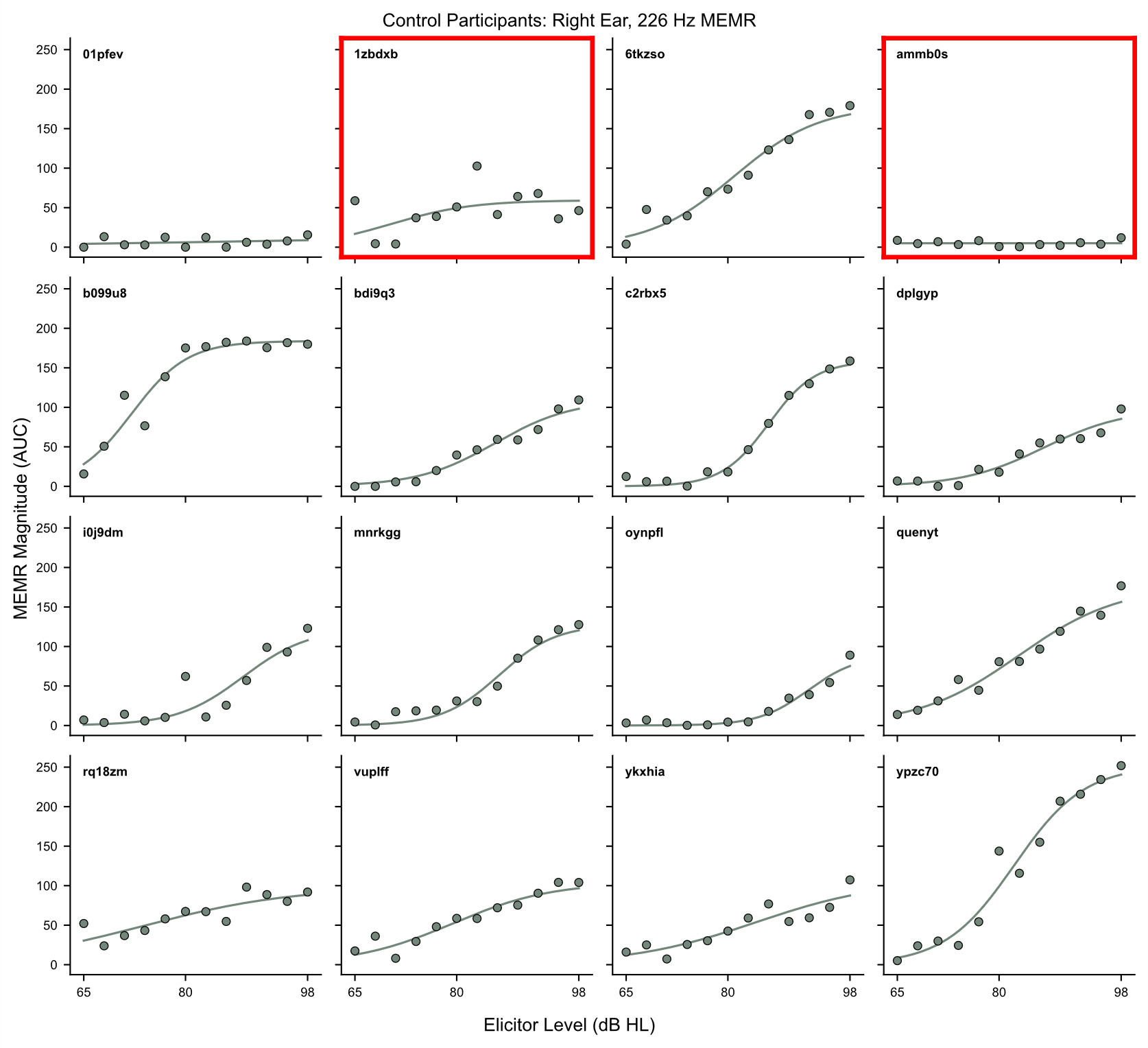

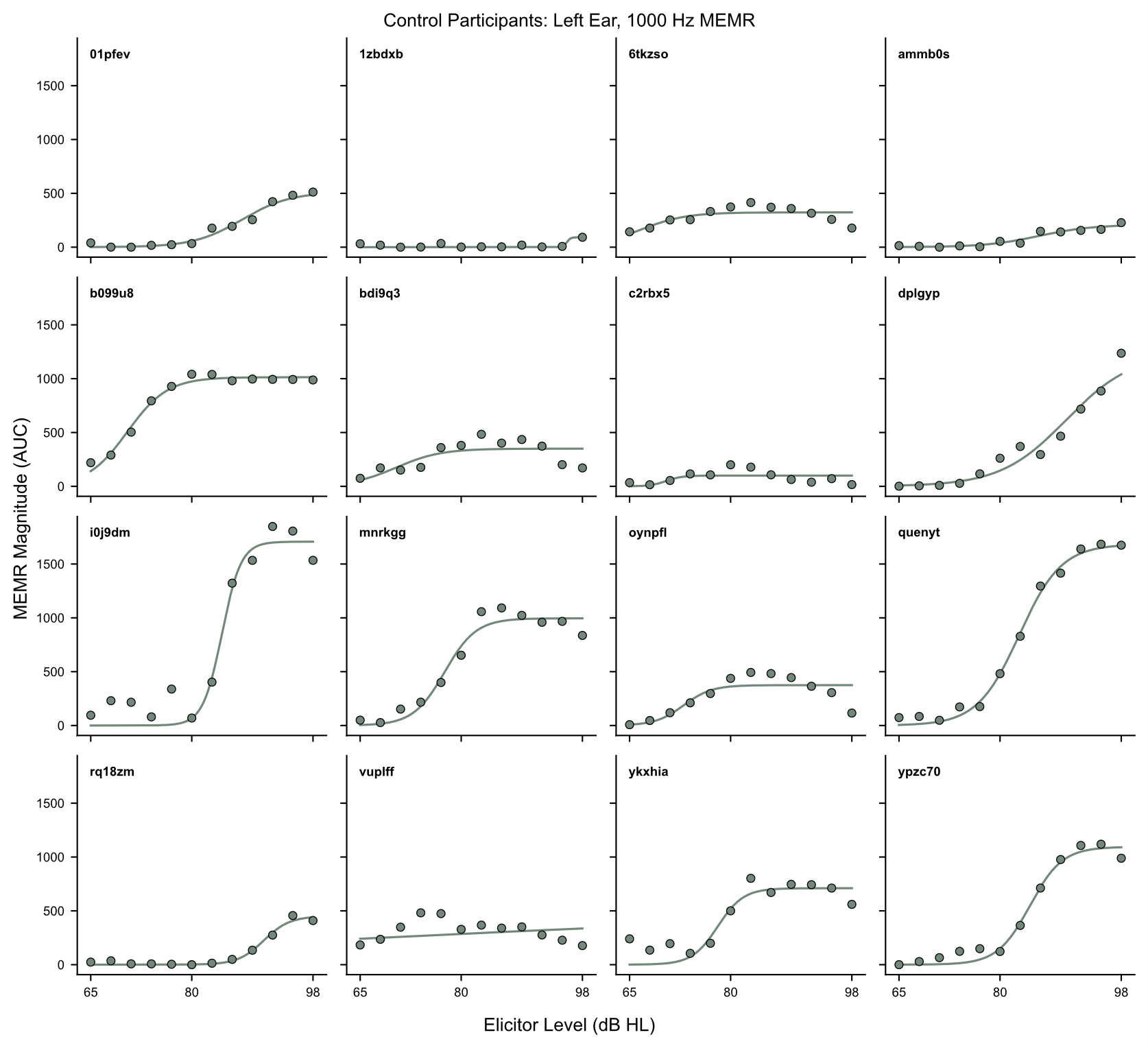

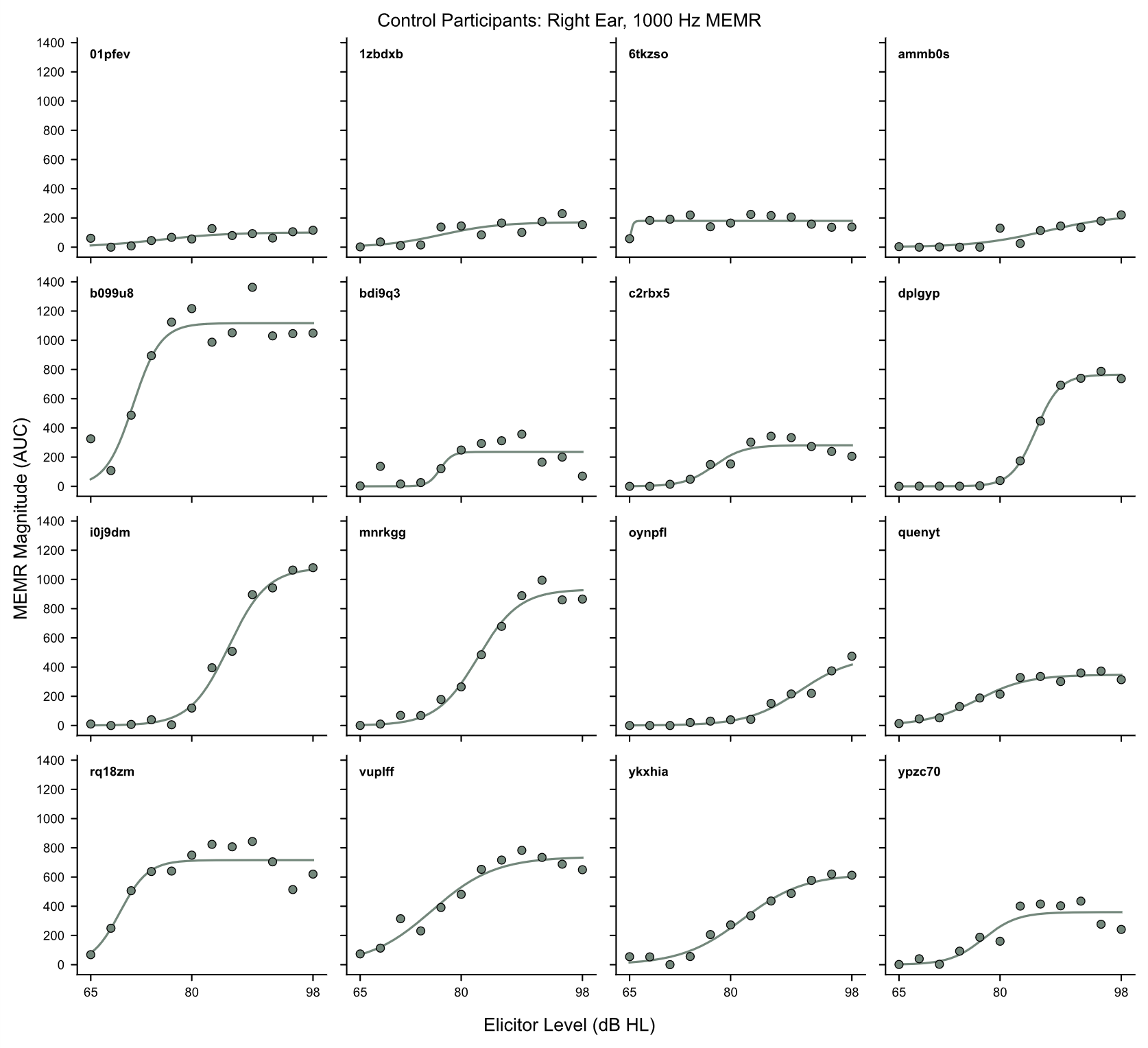

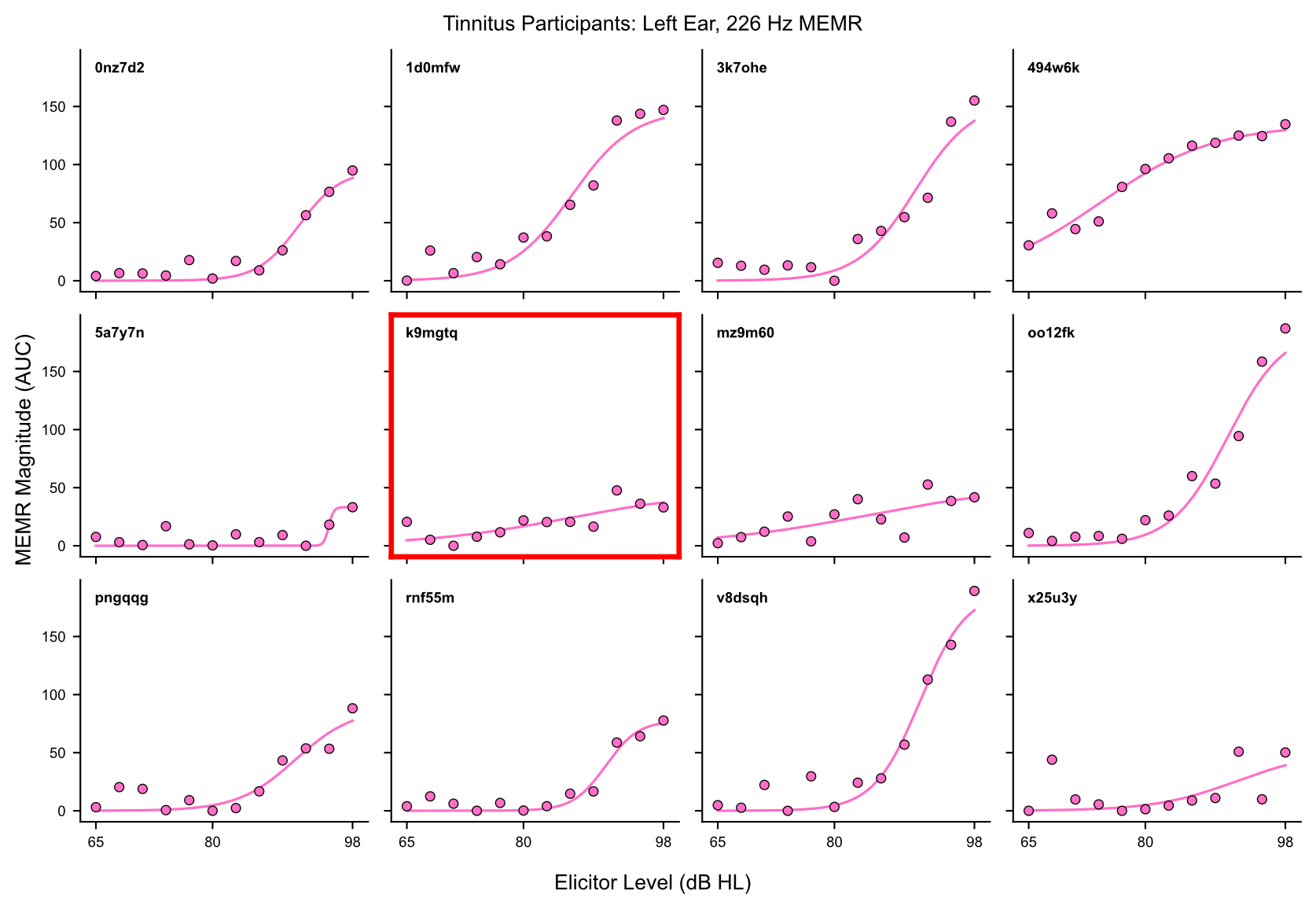

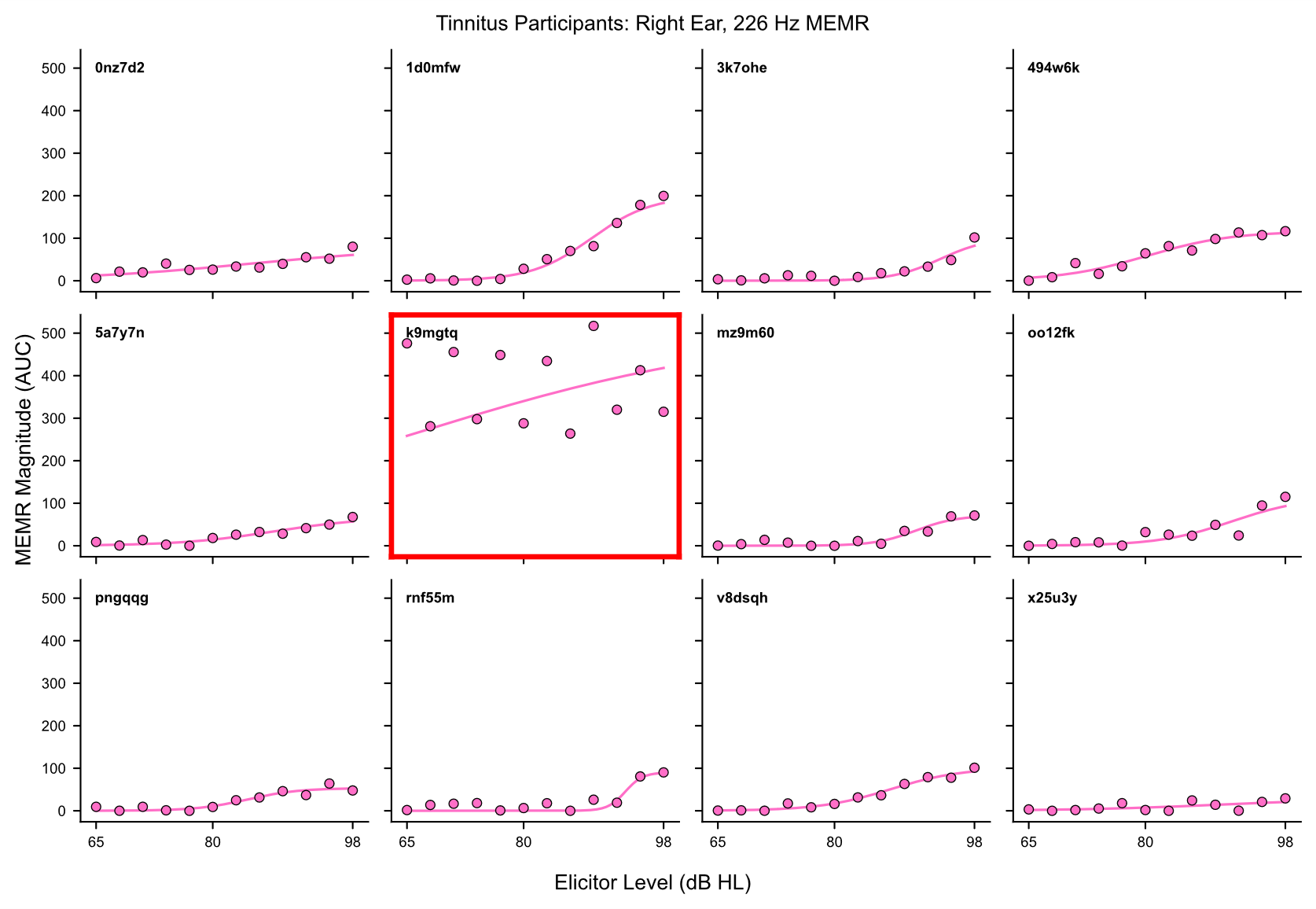

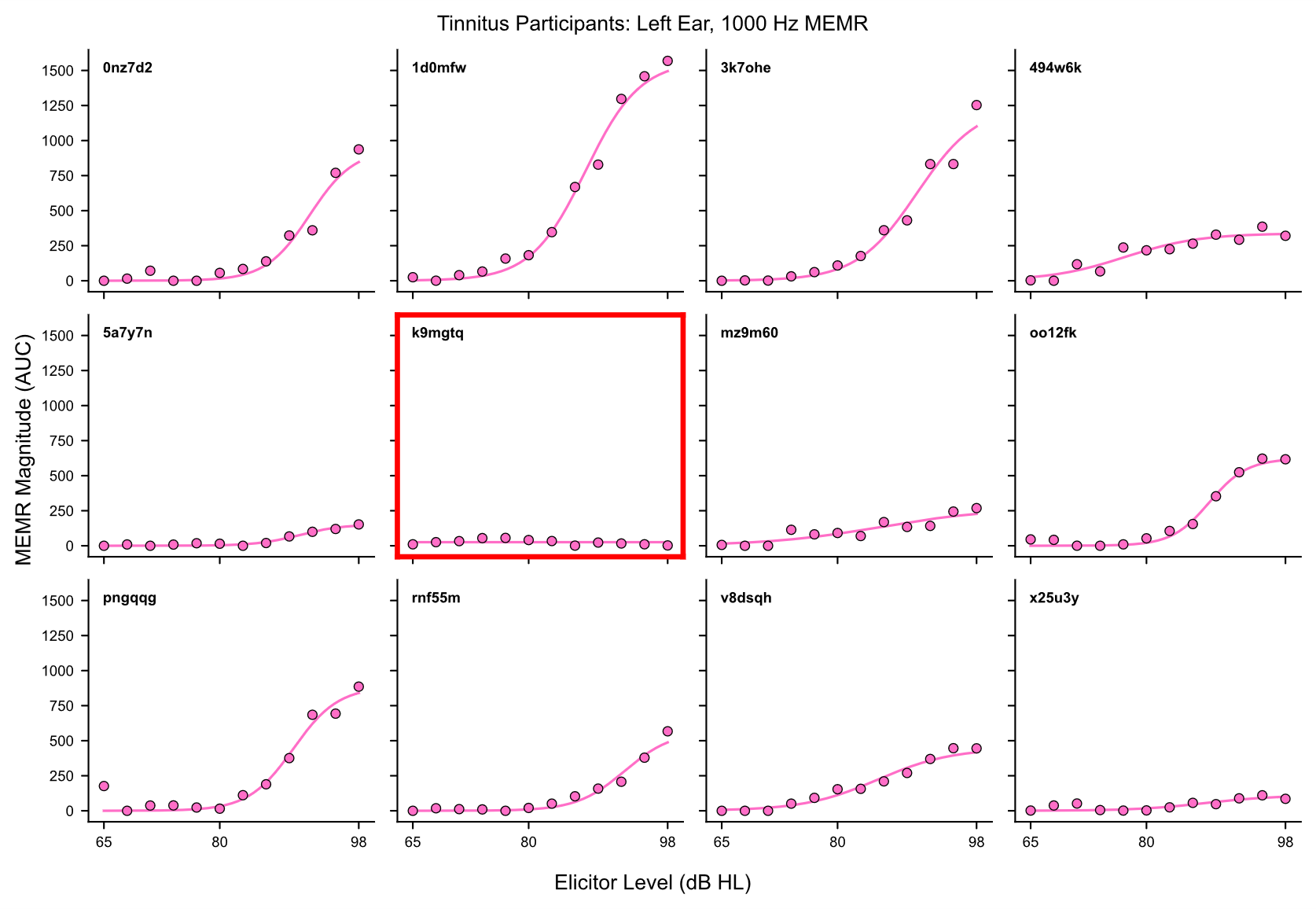

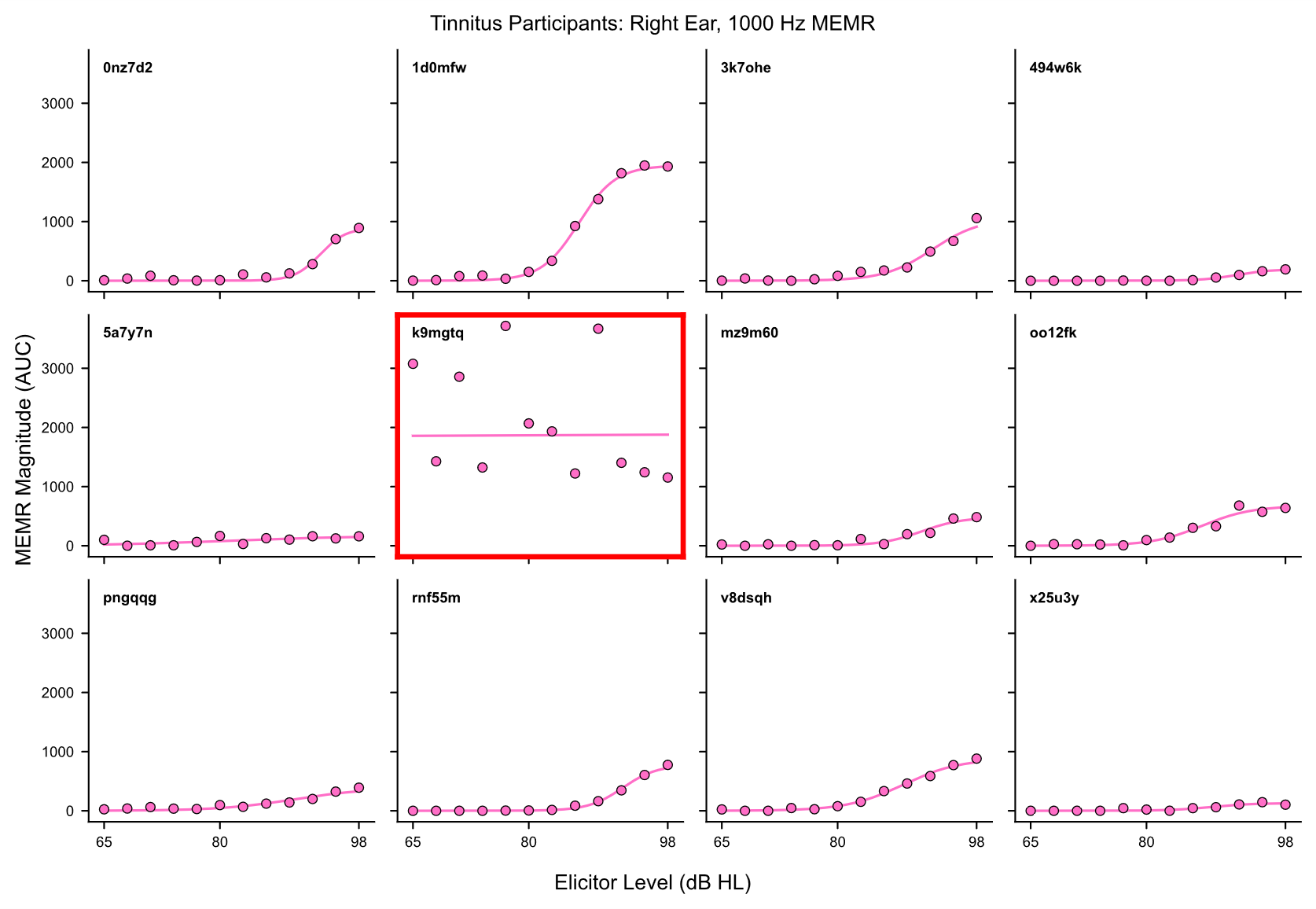

**SUPPLEMENTARY FIGURE S2.** Three-parameter sigmoid functions (lines) fitted to raw MEMR-by-level data (circles) for each probe tone and participant, with red boxes to highlight participants that were excluded due to unusual raw MEMR responses (data not shown) and/or poor sigmoid fits, as well as one extreme 226 Hz MEMR value (for participant i0j9dm), which was imputed via linear interpolation. Results for MEMR analyses were qualitatively the same with and without these changes.

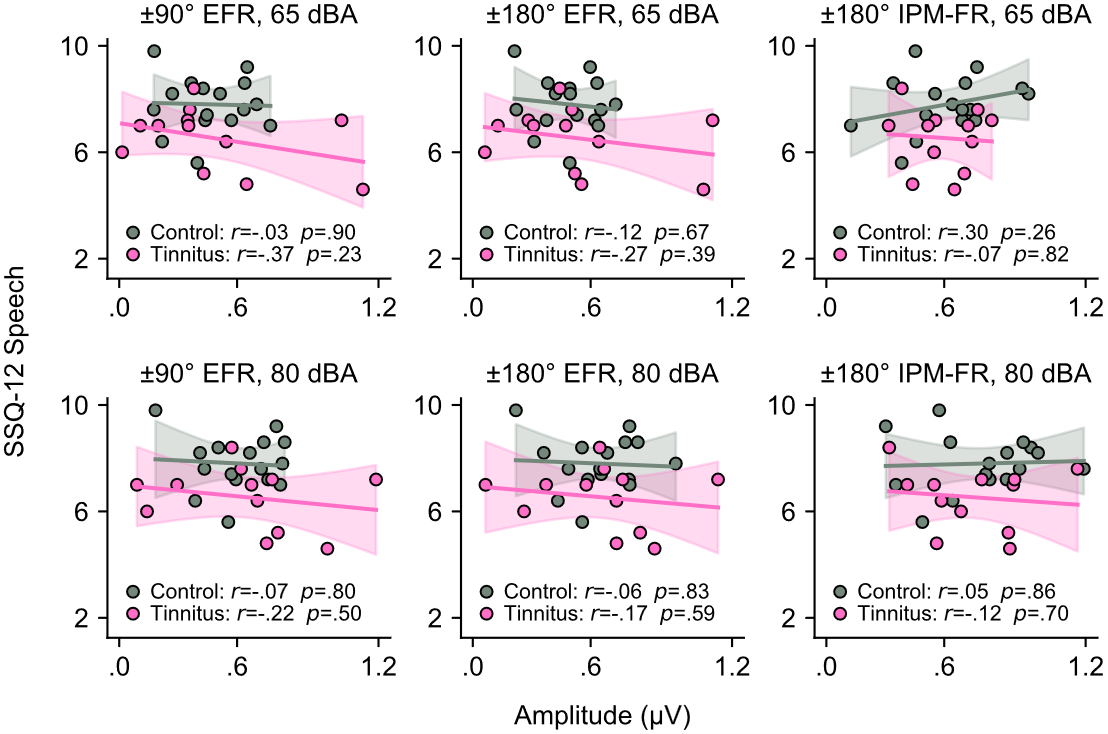

**SUPPLEMENTARY FIGURE S3.** Scatterplots depicting bivariate associations (not adjusted for age and sex) between SSQ-12 Speech scores and envelope following response (EFR) and 0°↔180° IPM-FR amplitudes, for each stimulus level. Shaded regions show the 95% confidence interval of the fitted line. There were no significant bivariate correlations, and linear mixed models (amplitude ~ group * speech * level + age + sex + (1 | subject)) did not show any significant main effects or interactions involving SSQ Speech or group on EFR or 0°↔180° IPM-FR amplitudes (see main text).
